# Synergism between a simple sugar and a small intrinsically disordered protein mitigate the lethal stresses of severe water loss

**DOI:** 10.1101/209163

**Authors:** Skylar X. Kim, Gamze Çamdere, Xuchen Hu, Douglas Koshland, Hugo Tapia

**Author notes:** Corresponding authors. (H.T.); (D.K.).

## Abstract

Anhydrobiotes are rare microbes, plants and animals that tolerate severe water loss. Understanding the molecular basis for their desiccation tolerance may provide novel insights into stress biology and critical tools for engineering drought-tolerant crops. Using the anhydrobiote, budding yeast, we show that trehalose and Hsp12, a small intrinsically disordered protein (sIDP) of the hydrophilin family, synergize to mitigate completely the inviability caused by the lethal stresses of desiccation. We show that these two molecules help to stabilize the activity and prevent aggregation of model proteins both *in vivo* and *in vitro*. We also identify a novel role for Hsp12 as a membrane remodeler, a protective feature not shared by another yeast hydrophilin, suggesting that sIDPs have distinct biological functions.

## INTRODUCTION

In the near future, global food security will be challenged by the effect of climate change on crop yields (Schwalm et al., 2017; Tirado et al., 2013). Potential solutions to more frequent drought may come from the study of drought-tolerant organisms. Indeed, diverse organisms, collectively named anhydrobiotes, can survive extreme water loss. By identifying and understanding the molecular mechanisms underlying desiccation tolerance, we hope to provide key new insights into stress biology and suggest fruitful avenues for the engineering of drought-tolerant crops.

For almost a half a century, scientists have known that almost all anhydrobiotes have high levels of the disaccharide trehalose and hydrophilins, which are small, hydrophilic, and intrinsically disordered proteins (sIDPs) (Crowe et al., 1992; Crowe, 1971; Potts, 2001). Demonstrating a functional significance for this striking correlation eluded researches until very recently. A requirement for trehalose production in desiccation tolerance was shown in genetic studies of two anhydrobiotes, yeast and nematodes (Erkut et al., 2011; Tapia and Koshland, 2014). However, their requirement for trehalose was not identical; nematodes that were unable to synthesize were sensitive to desiccation under all conditions while yeast that were unable to synthesize trehalose were sensitive to long-term, but not short-term, desiccation (Erkut et al., 2011; Tapia and Koshland, 2014).

Similarly, recent studies have also revealed disparate requirements for hydrophilins in desiccation tolerance. In tardigrades, RNAi knock-down of individual hydrophilins caused desiccation sensitivity (Boothby et al., 2017). In contrast, in yeast, mutants lacking a subset of either individual or multiple hydrophilins display normal tolerance to desiccation (Calahan et al., 2011). The different requirements of both hydrophilins and trehalose in different anhydrobiotes suggest that overlapping activities, potentially of trehalose, hydrophilins, or additional, as yet unidentified, stress effectors mitigate desiccation tolerance. Therefore, a major unanswered question in the field, is what stress effectors are necessary and sufficient for desiccation tolerance?

The identification of this set of stress effectors might provide a foundation to understand the stresses imposed by desiccation. In principle, the lack of intracellular water could readily lead to changes in proteostasis, membrane permeability, and/or osmolarity. Anyone or all of these changes might cause lethality. Characterizing the molecular activities of the stress effectors that promote desiccation tolerance could identify the critical/lethal stresses of desiccation.

Recent studies in budding yeast suggest that it is an excellent model anhydrobiote to identify the stress effectors required for desiccation tolerance and to characterize their molecular activities (Calahan et al., 2011; Tapia and Koshland, 2014; Tapia et al., 2015; Welch et al., 2013). Importantly, budding yeast cells display conditional desiccation tolerance. In stationary phase, 20-40% of cells survive desiccation. However, when dividing exponentially, only approximately 0.001% of cells survive desiccation (Calahan et al., 2011). This conditional tolerance provides a powerful genetic tool to assess the contributions of different factors to desiccation tolerance.

The loss of desiccation tolerance, in mutant yeast cells grown to stationary-phase, has been used to identify genes necessary for desiccation tolerance (Calahan et al., 2011). Examination of the yeast, non-essential gene deletion collection, for mutants unable to tolerate short-term desiccation tolerance, identified deletions in genes that block respiration (Calahan et al., 2011). While identifying an essential process such as respiration as being important for desiccation tolerance, this screen did not inform on potential stress effectors because changes in respiration alters the expression of many hundreds of genes, including many potential stress effectors. Strikingly, this genetic screen failed to identify deletions in non-regulatory genes. The absence of these deletions supported the notion that stress effectors with overlapping functions in desiccation tolerance must exist, precluding their identification by a single gene deletion.

An increase in desiccation tolerance of exponentially dividing yeast cells provides a measure of the sufficiency of individual factors to confer desiccation tolerance. Indeed, elevated intracellular trehalose or expression of several individual tardigrade sIDPs increased the survival of exponentially-dividing yeast after desiccation by 1000-fold (Boothby et al., 2017; Tapia et al., 2015). These results showed that trehalose and sIDPs could act as potential stress effectors in the same organism. However, the level of desiccation tolerance in cells with elevated trehalose or the tardigrade sIDPs was still significantly less than the tolerance in stationary-phase cells, further reinforcing the notion that tolerance likely requires multiple effectors working in concert.

We sought to identify such effectors and to characterize their molecular functions. Here, using our genetic assays with stationary-phase and exponentially growing yeast cells, we demonstrate that trehalose and Hsp12, a yeast hydrophilin, synergize to completely alleviate all of the lethal damage that occurs due to severe water loss. We provide evidence that these two small molecules counteract the misfolding and aggregation of protein reporters that occurs upon desiccation both *in vitro* and *in vivo*. We identify a novel role for Hsp12 as a membrane remodeler, an activity likely required to provide desiccation tolerance. Finally, our data suggests that despite sharing many of the same properties (high glycine content, small size, high hydrophilicity, disordered secondary structure), yeast hydrophilins clearly have distinct functions. We discuss the implications of these findings for both stress biology and potential translational applications.

## RESULTS

### Trehalose and Hsp12 are necessary and sufficient for desiccation tolerance

We reasoned that the previous screen of the non-essential yeast deletion collection for sensitivity to short-term desiccation sensitivity was unable to identify individual stress effectors because likely more than one stress effector needed to be inactivated to compromise tolerance. At least one of these factors was likely trehalose, given its role in long-term desiccation tolerance. Therefore, we started with a base strain that was unable to synthesize trehalose (*tps1∆*) and introduced into it deletions for the remaining ~5,000 non-essential genes (Tong et al., 2001). These 5000 double mutants were grown to stationary phase and assayed for survival after short-term and long-term desiccation. We identified three groups of double mutants with distinct phenotypes. The first group of strains were inviable when subjected to short-term desiccation (Table S1A, S1-2). The second group of strains were inviable even in the absence of stress (Table S1-1). The third group suppressed the long-term desiccation sensitivity of a *tps1∆* (Table S1-3). While these last two groups identify many potentially interesting genes, we focused on the first group; the gene deletions in this group potentially inactivated candidate stress effectors alongside trehalose that were needed for short-term desiccation tolerance.

One candidate gene from our desiccation sensitive group was *HSP12*, which encodes one of 12 previously identified yeast hydrophilins and is the most highly expressed member of the family (Garay-Arroyo et al., 2010). To investigate the potential synergism between Hsp12 and trehalose further, we compared quantitatively short-and long-term desiccation tolerance of stationary cells that were wild type, *tps1∆, hsp12∆*, or *tps1∆ hsp12∆*. After a 30-day desiccation period, the viability of stationary wild type cells decreased only two-fold (Figure 1A). Although the viability of *tps1∆* cells was similar to that of wild type cells after 2 days of desiccation, after the 30-day desiccation period, *tps1∆* cell viability dropped more than 100-fold (Figure 1A). This result corroborated our previous evidence demonstrating a requirement of trehalose for long-term but not short-term desiccation tolerance (Tapia and Koshland, 2014). While *hsp12∆* cells exhibited wild type levels of desiccation tolerance after both short-and long-term desiccation, the *tps1∆ hsp12∆* cells displayed a 100-fold drop in viability after only 2 days of drying. Furthermore, after 30 days of desiccation, no viable *tps1∆hsp12∆* cells remained (Figure 1A). These results revealed that trehalose and Hsp12 act cooperatively to promote tolerance to both short-and long-term desiccation. Furthermore, although *tps1∆* and *hsp12∆* had both been shown to confer elevated sensitivity to heat stress, the *tps1∆ hsp12∆* double mutant was no more heat-sensitive than either single mutant alone (Figure S1) (Gibney et al., 2015; Welker et al., 2010). Thus, the protective synergism we observe between trehalose and Hsp12 is specific to desiccation stress.

We had shown previously that exponentially growing cells with increased intracellular trehalose showed a 1000-fold increase in desiccation tolerance (Tapia et al., 2015). Hence, we tested whether exponentially dividing cells that expressed high-level expression of Hsp12 would also exhibit increased desiccation tolerance. Normally, Hsp12 is not highly expressed in unstressed dividing cells. Therefore, we expressed Hsp12 under a strong constitutive promoter to drive expression in dividing cells to a level similar to stationary cells. These dividing cells exhibited a 1000-fold increase in the desiccation, comparable to the effect of increased intracellular trehalose (Figure 1B). Thus, Hsp12, like trehalose, had a significant ability to mitigate one or more of the lethal stresses imposed by desiccation.

**Figure 1.**
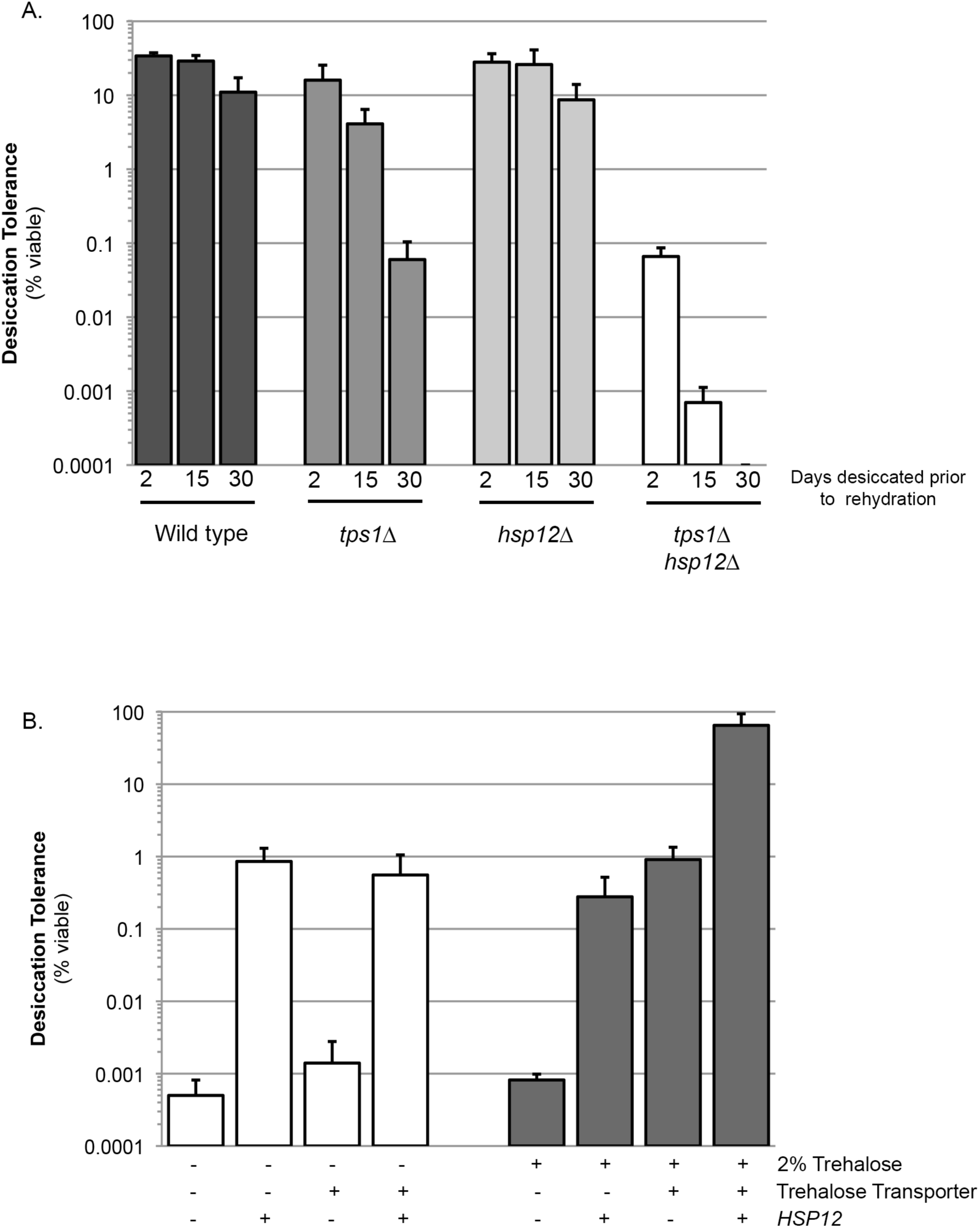
Trehalose and Hsp12 are necessary and sufficient for desiccation tolerance. (**A**) Yeast cells were grown to saturation (5 days), air-dried for 2, 15, or 30 days at 23 °C, 60% relative humidity (RH), then rehydrated and assessed for viability by counting colony forming units (CFU). Desiccation tolerance of wild type (BY4742), HTY104 (*tps1∆*), HTY11 (*hsp12∆*) or HTY176 (*tps1∆hsp12∆*) cells. (**B**) Yeast cells were grown to mid-exponential phase (OD < 0.5) in selective media (SC-His+Dextrose). Cells were then transferred to either selective media (SC-His+Dextrose, 0% trehalose) or selective media with trehalose (SC-His+Dextrose, 2% trehalose) for 1 h. Cells were collected, washed, and air dried for 2 d at 23 °C, 60% relative humidity (RH), then rehydrated and assessed for viability by counting colony forming units (CFU). Log-phase experiments performed in strain HTY121 (*nth1∆* (trehalase delete)), additionally: +/-Trehalose transporter (*AGT1* expressed constitutively (*TDH3pr-AGT1)*) and +/-*HSP12* (Hsp12 expressed constitutively from a plasmid (HTP126), strains not over-expressing *HSP12* carry an Empty Vector plasmid (HTP64).

Next, we looked for potential synergy between trehalose and Hsp12 by examining the desiccation tolerance of exponentially dividing yeast cells containing high levels of both protectants. The desiccation tolerance of these cells was ~60-fold greater than cells that expressed only Hsp12 or trehalose. Furthermore, the absolute amount of tolerance (65-80% survival) was even greater than the tolerance of wild type cells in stationary phase (20-40% survival), the highest tolerance previously reported for yeast cells (Figure 1B) (Calahan et al., 2011). Our findings suggest that the presence of these two stress effectors together are sufficient to counter all the lethal effects of desiccation and subsequent rehydration on exponentially dividing yeast.

### Trehalose and Hsp12 modulate proteostasis *in vivo* and *in vitro*

Having previously established that trehalose helped prevent desiccation-induced proteotoxicity *in vivo* (Tapia and Koshland, 2014), we tested the *in vivo* role of Hsp12, with and without trehalose to modulate proteostasis. We first utilized the inactivation of firefly luciferase as a proxy for misfolding and aggregation *in vivo.* As a control, we expressed luciferase in exponentially dividing wild-type cells that are expressing neither trehalose nor Hsp12. We also expressed luciferase in exponentially dividing cells with high levels of Hsp12, trehalose or both protectants together. After desiccation, luminescence in wild type cells was reduced one hundred thousand-fold (Figure 2A). Thus, luciferase activity was extremely sensitive to desiccation. Cells with elevated trehalose retained a four-fold increase in luminescence; while cells with Hsp12 alone failed to retain any increase in luminescence after drying. However, the luminescence of cells that contained both elevated trehalose and elevated Hsp12 was 14-fold greater than wild type and 4-fold greater than cells with just elevated trehalose (Figure 2A). These results demonstrate that Hsp12 can enhance the small, but reproducible, ability of trehalose to prevent luciferase inactivation.

**Figure 2.**
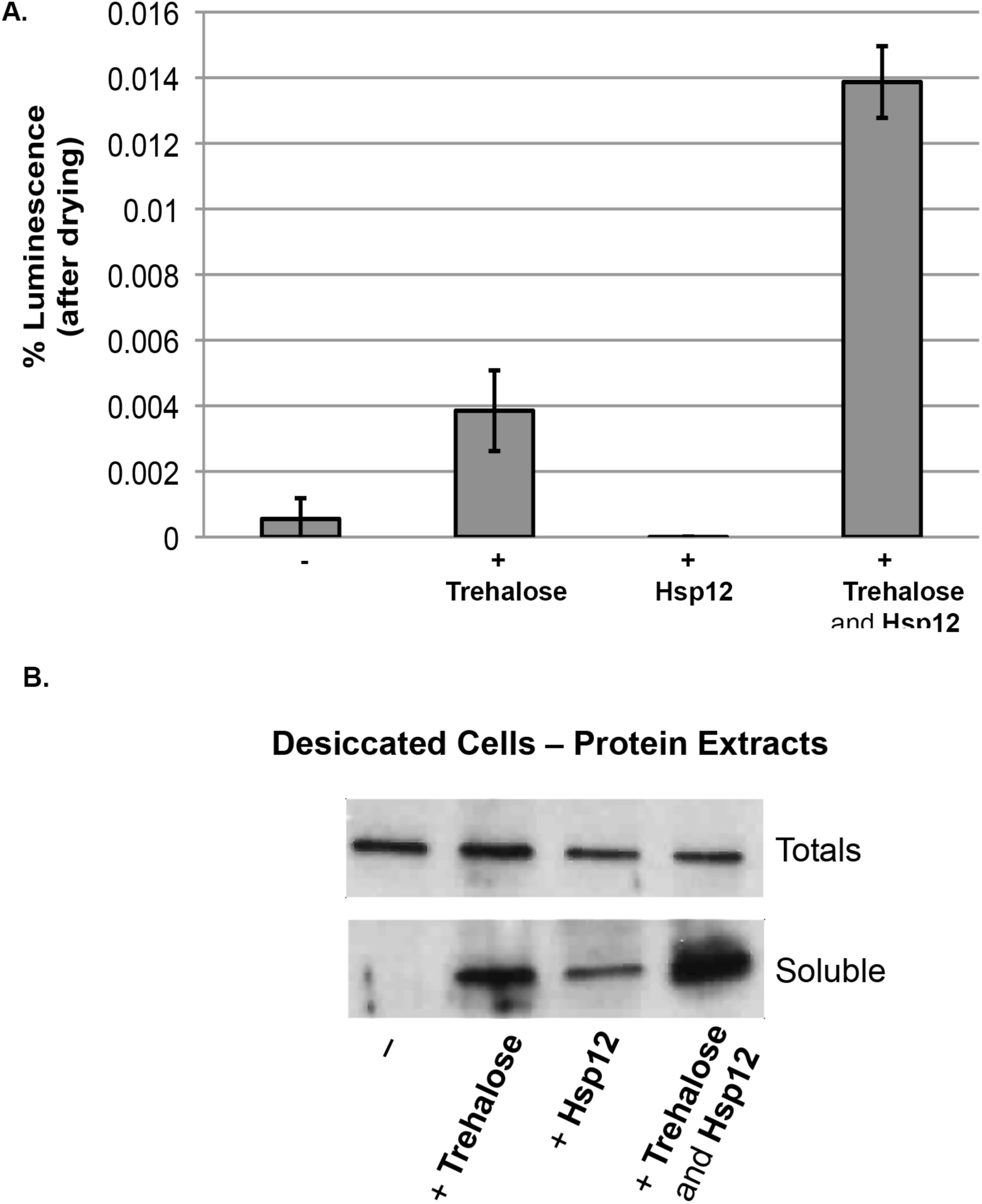
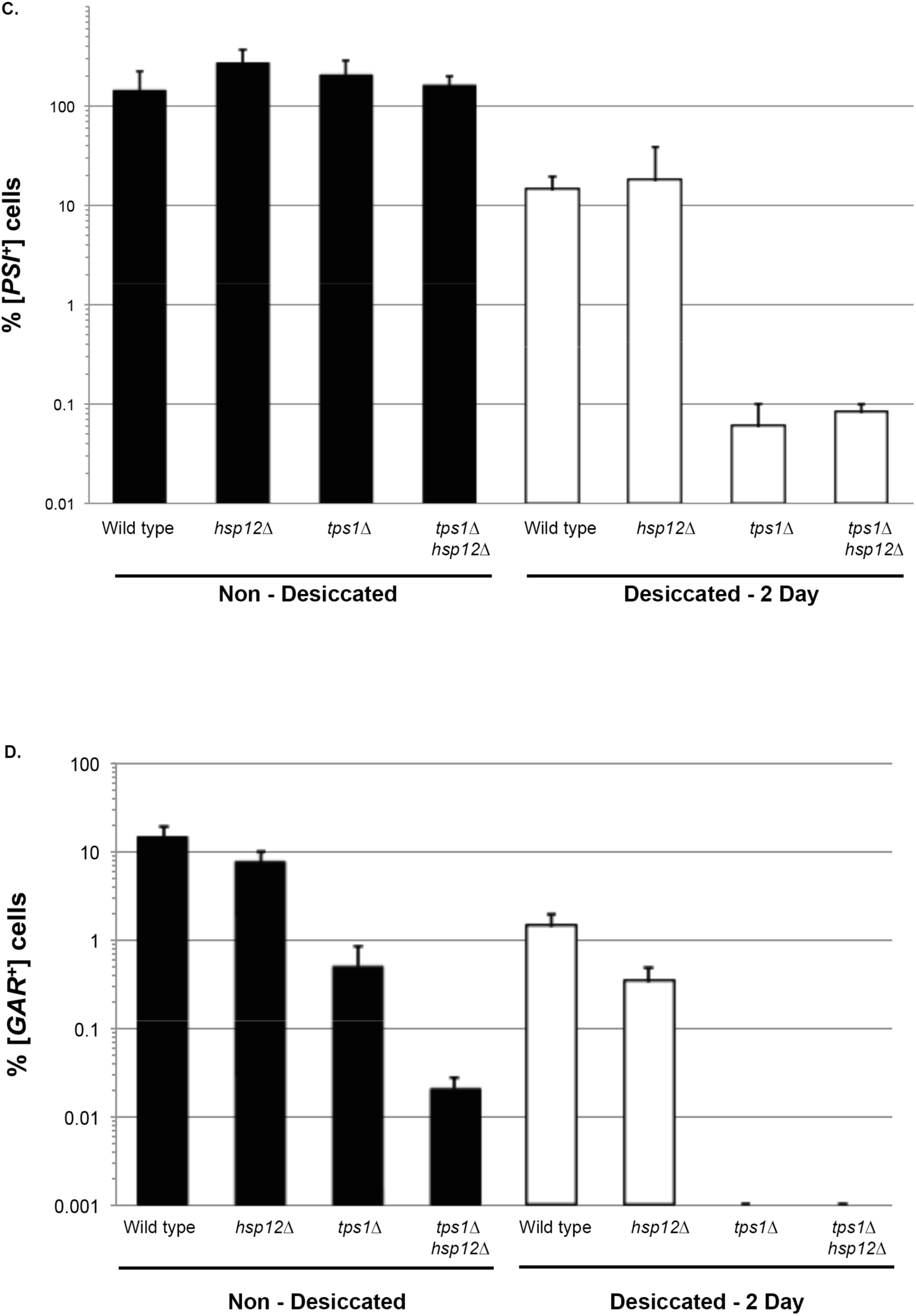
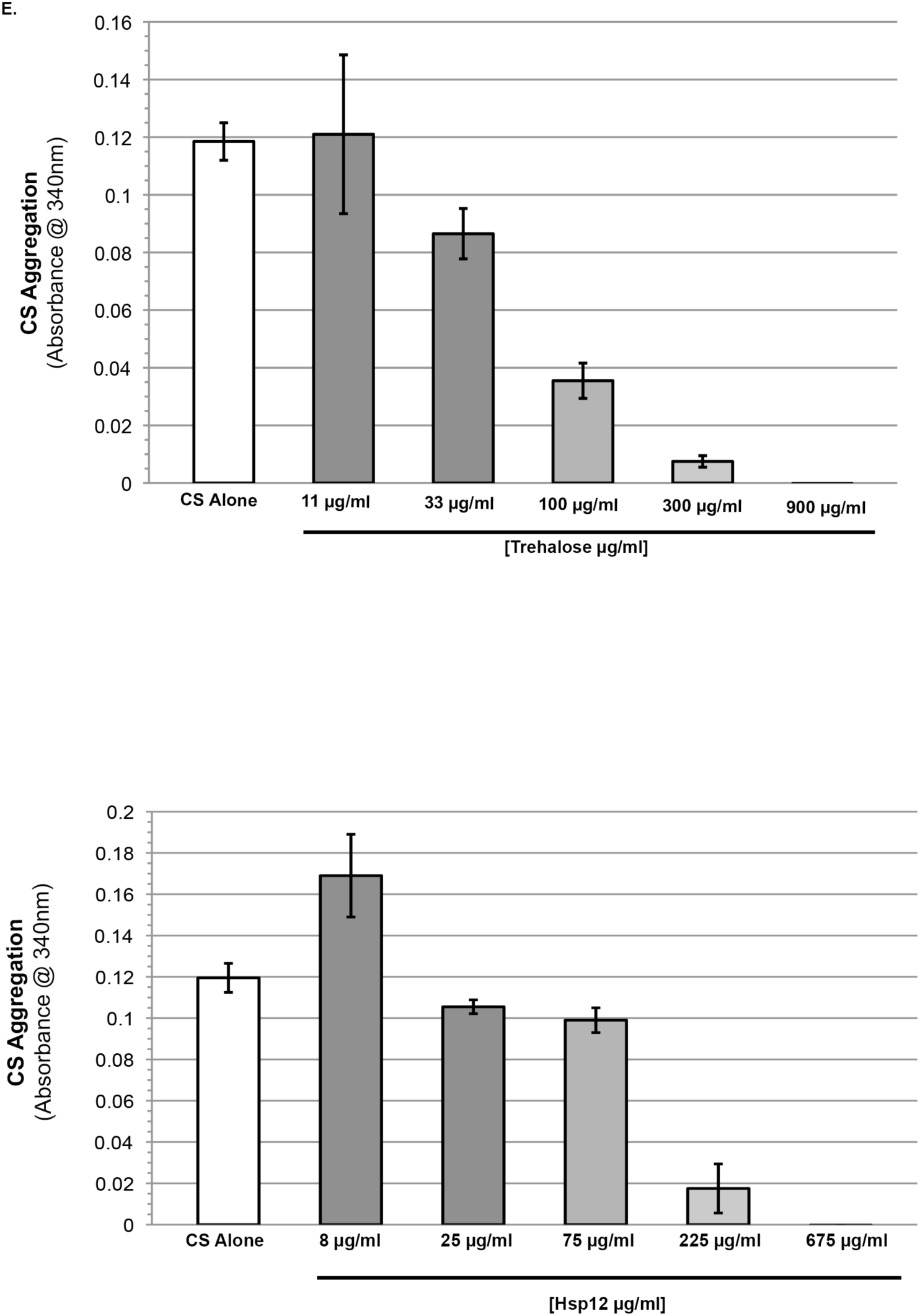
Trehalose and Hsp12 stabilize activity and prevent aggregation of *in vivo* and *in vivo* model substrates. (**A-B**) Luciferase activity and aggregation as a reading of proteostasis of logarithmically growing and desiccated cells. (**A**) Yeast cells were transformed with a temperature-sensitive firefly luciferase protein expressed from the constitutive (GPD) promoter (HTP111) or empty vector (HTP82). Strains were grown to mid-log phase in selective media (SC-Ura-His+Dextrose) and then transferred to selective media with trehalose (SC-Ura-His+Dextrose, 2% trehalose) for 1 h. Desiccated samples were air-dried for 2 days followed by rehydration in SC-Ura-His + cycloheximide (10 µl/ml, to block new FFL protein synthesis). Luciferase activity 682 was measured in *vivo* by addition of 0.5 mM D-Luciferin to equal number of intact cells. Light emission was measured immediately with a luminometer. Percentage of activity was normalized to WT saturated ‘‘wet’’ sample. Luciferase experiments performed in strain HTY163 (*nth1∆* (trehalase delete)), additionally: +/-Trehalose transporter (*TDH3pr-AGT1)* and +/-Hps12 (HSP*12* expressed constitutively from a plasmid (HTP126), strains not over-expressing *HSP12* carry an Empty Vector plasmid (HTP64). (**B**) Cells were prepared as in A. Total cellular protein was extracted, and aggregated proteins were sedimented by high-speed centrifugation. The amounts of luciferase in the total protein fraction (total) and in high-speed supernatants (soluble) were determined by SDS-PAGE and followed by immunoblotting with antiserum recognizing luciferase. (**C-D)** Prion propagation as a measure of *in vivo* protein propagation after desiccation. (**C**) [*PSI*^+^] prion propagation before, and after desiccation. To assess [*PSI*^+^] prion state, cells were plated on media lacking adenine (SC-ADE) compared to non-dried controls. (**D**) [*GAR*^+^] prion propagation before, and after desiccation. To assess [*GAR*^+^] prion state, cells were plated on YP media with 2% glycerol and 0.05% glucosamine compared to non-dried controls. Strain information available on strain Table S2. (**E**) Desiccation-induced aggregation of citrate synthase in the presence of varying concentrations of trehalose or Hsp12. Absorbance at 340 nm. Citrate Synthase (0.12 mg) and increasing concentrations of trehalose, or Hsp12. Absorbance measured after drying at 36°C for 24 hours followed by hours at 23°C, then rehydration in water. Measurements after drying compared to 701 measurements before drying.

To complement the luminescence assay, we examined desiccation-induced luciferase aggregation. Lysates were prepared from our panel of desiccated exponentially dividing cells immediately after rehydration. The lysates were subjected to centrifugation to separate soluble luciferase from insoluble luciferase that had aggregated with itself or other insoluble proteins. For all strains tested, equivalent amounts of total luciferase were present in cell lysates (Figure 2B). In the wild type strain lacking trehalose and Hsp12, no luciferase was detected in the soluble fraction (Figure 2B). Increased intracellular trehalose or Hsp12 significantly increased luciferase solubility. Despite increasing solubility, Hsp12 failed to rescue any luciferase activity (Figure 2A). Thus Hsp12 was more potent at blocking luciferase aggregation than it was at preventing its denaturation. We conclude that desiccation, a naturally-occurring environmental condition, potentiates denaturation and aggregation of luciferase, and these proteotoxic effects can be mitigated *in vivo* by either trehalose, Hsp12, but more pronouncedly by a combination of the two.

We next examined prion propagation as a second independent measure of the *in vivo* role of trehalose and Hsp12 in proteostasis. Prion propagation provides two measures of proteostasis. It requires stabilization of an altered protein state and a balance of oligomerization, enough to transmit the altered protein state to other proteins but not too much to prevent its transmission to daughter cells during division. Previously, we used [*PSI*^+^], a cytoplasmic prion or [*GAR*^+^], a membrane-associated prion, to show that trehalose prevents hyper-aggregation during desiccation of stationary yeast cells (Tapia and Koshland, 2014). With this precedent, we constructed wild type, *tps1∆*, *hsp12∆*, or *tps1∆ hsp12∆* strains with either [*PSI*^+^] or [*GAR*^+^]. We grew aqueous cultures of these strains to stationary phase and determined the percentage of cells that contained prions by growing on selective media specific to the presence of the different prions. The cells in these cultures were also subjected to two days of desiccation and then assessed for their ability to propagate the prions to their rehydrated descendants.

Prior to desiccation, the percentage of stationary cells with [*PSI^+^*] was statistically indistinguishable in wild type, *hsp12∆*, *tps1∆*, or *tps1∆ hsp12∆* cultures (Figure 2C). [*PSI*^+^] propagation was dramatically reduced in the descendants of desiccated *tps1∆* cells (Figure 2C) as expected from our previous study (Tapia and Koshland, 2014). In contrast, the lack of Hsp12 (*hsp12∆*) alone did not affect the propagation of [*PSI*^+^] after desiccation more than wild type cells, nor did it exacerbate the loss of [*PSI*^+^] propagation in cells unable to make trehalose (*tps1∆ hsp12∆*) (Figure 2C). These results suggest that trehalose is more effective than Hsp12 in preventing aggregation of this cytoplasmic prion.

In contrast, trehalose and Hsp12 exhibited a striking synergism in the propagation of [*GAR*^+^], a membrane prion. In *tps1∆* cells, the percentage of [*GAR*^+^] cells was ~10-fold lower than wild type after desiccation (Figure 2D) as expected from our previous study (Tapia and Koshland, 2014). The percentage [*GAR*^+^] cells did not change significantly in *hsp12∆* cultures before or after transient desiccation (Figure 2D). In *tps1∆ hsp12∆* cultures, the percentage of [*GAR*^+^] cells was undetectable after desiccation, as expected given that *tps1∆* cells were unable to propagate [*GAR*^+^] through desiccation. However, even in the absence of desiccation, the percentage of [*GAR*^+^] cells in *tps1∆ hsp12∆* cultures reduced almost 500-fold compared to wild type cells (Figure 2D). Thus, these two small effectors act synergistically to modulate the proteostasis of the [*GAR*^+^] membrane prion even in the presence of water. This synergism also may occur during desiccation, but was masked by the severe effect of the loss of trehalose alone.

To test whether the impact of trehalose and Hsp12 on proteostasis was direct, we examined their effect *in vitro* on the desiccation-induced aggregation of citrate synthase (CS). We desiccated solutions of CS alone, or with trehalose, Hsp12 or both of the stress effectors together. These samples were rehydrated, and CS proteostasis was assessed by assaying its aggregation and its enzymatic activity.

Desiccation of CS caused a greater than 20-fold increase in its aggregation and a loss of approximately 50% of its enzymatic activity (Figure 2E, S2A). The addition of increasing amounts of trehalose or Hsp12 before desiccation decreased the desiccation-induced aggregation of CS six to twelve-fold respectively. (Figure 2E). Both trehalose and Hsp12 also restored nearly all CS enzymatic activity in a concentration-dependent manner (Figure S2A). Limited synergism was observed between trehalose and Hsp12 against desiccation-induced damage of this protein substrate (Figure S2B-C). The range of trehalose concentration that impacted CS aggregation was very similar to the range of intracellular trehalose required to promote desiccation tolerance *in vivo* (Tapia et al., 2015). Significant solubilization of CS occurred at a 1:1 ratio of Hsp12 to CS (Figure 2E, 225 µg/ml). This equal stoichiometry suggests that the proteostasis function of Hsp12 is not enzymatic, consistent with its high expression during stress *in vivo*. These results suggest that both trehalose and Hsp12 have independent proteostasis activities *in vitro* that may contribute to their protective *in vivo* activities.

### Hsp12 remodels lipid vesicles

The synergistic function of trehalose and Hsp12 in [*GAR*^+^] propagation could be due to their direct effect on membrane proteins and/or an indirect effect on overall membrane structure and integrity. Intriguingly, prior studies indicate that trehalose stabilizes membranes *in vitro* and this stabilization may prevent disruptive phase transitions that occur upon rehydration (Crowe et al., 1984; 1987; Erkut et al., 2011; Leslie et al., 2002; Penkov et al., 2010; Tang et al., 2007). Hsp12 has also been reported to increase membrane stability (Welker et al., 2010). A potential membrane function for Hsp12 was also suggested by the fact that, although Hsp12 does not exhibit any detectable secondary structure in solution, it acquires an alpha-helical conformation in the presence of certain lipids (Welker et al., 2010). Secondary structure prediction and NMR data identify four alpha-helical regions in Hsp12, with each of the three larger helices having the features of amphipathic helices (Singarapu et al., 2011; Welker et al., 2010). Numerous membrane-remodeling proteins use amphipathic helices, embedding the hydrophobic face partway into the bilayer to shape membranes (Boucrot et al., 2012; Meinecke et al., 2013). Given these considerations, we hypothesized that Hsp12 might have membrane remodeling activities.

One readout of membrane remodeling activity is the capacity to vesiculate large liposomes into nanovesicles (Boucrot et al., 2012). For example, proteins known to be involved in endocytic membrane trafficking lead to the vesiculation of liposomes (Boucrot et al., 2012). Therefore, we tested whether addition of Hsp12 to dimyristoylphosphatidylglycerol (DMPG) liposomes would generate nanovesicles with bound Hsp12. DMPG liposomes incubated in the absence of Hsp12 sedimented into the pellet fraction due to their large size (Figure 3A). By contrast, when incubated in the presence of Hsp12, DMPG appeared in the supernatant fraction (Figure 3A). To assess vesiculation directly, we examined these same samples by electron microscopy. Prior to the addition of Hsp12, liposomes exhibited expected shapes and sizes. After the addition of Hsp12, very few intact liposomes were observed, replaced by membrane remnants (Figure 3B). Trehalose alone did not have any vesiculation activity nor did it inhibit the effect of Hsp12 on the liposomes. These results show that Hsp12, but not trehalose, has membrane-remodeling activities *in vitro*. Our findings suggest that their synergistic impact on [*GAR*^+^] propagation may result from different activities.

**Figure 3.**
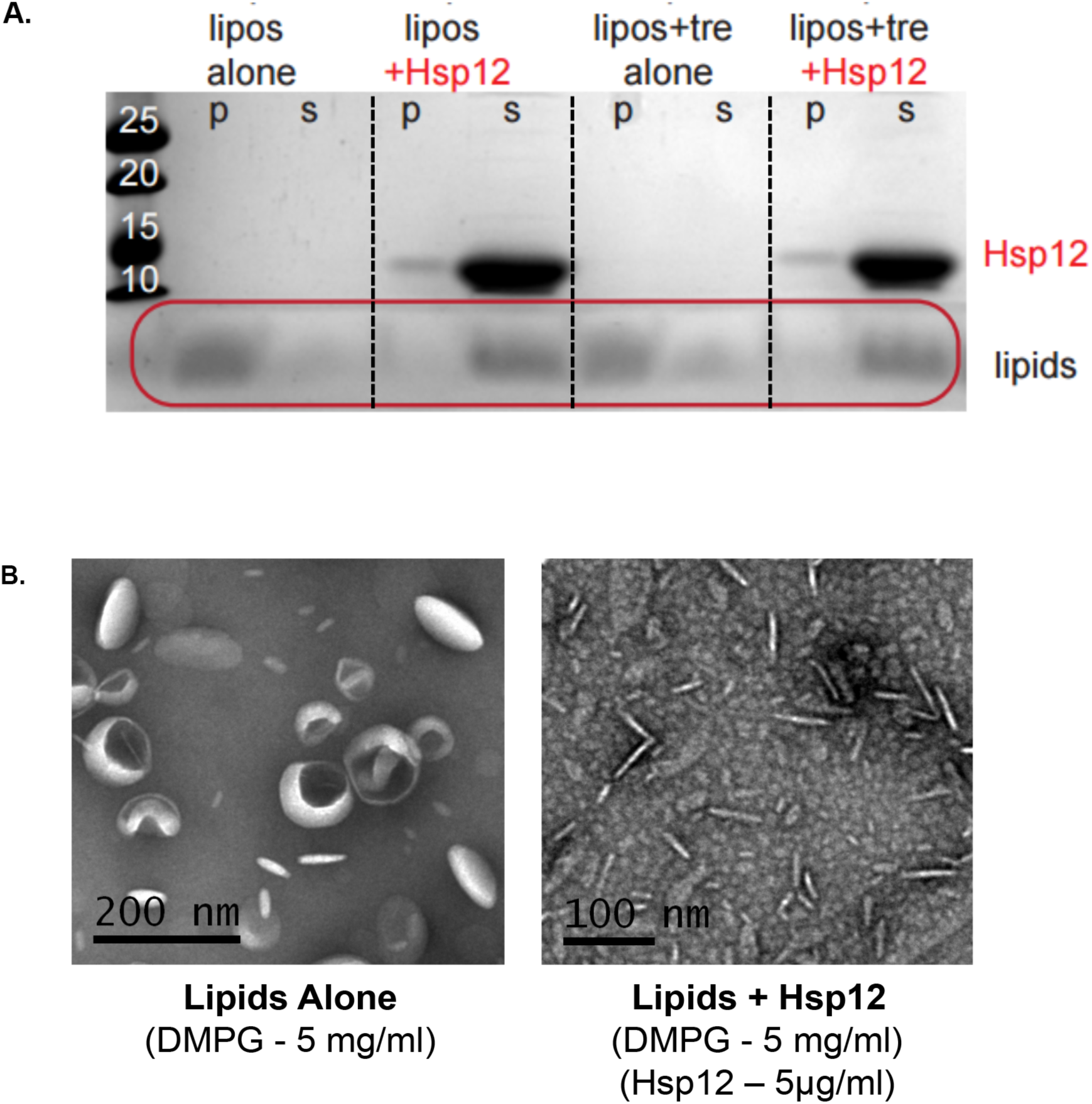
Hsp12 causes membrane remodeling. (**A**) DMPG liposomes (5 mg/ml) where incubated for 1 hour at room temperature in the 714 presence or absence of Hsp12 (25 ug/ml) and/or in the presence or absence of trehalose (2% final). Pellet (p) and supernatant (s) fractions where separated by high-speed centrifugation, and lipid distribution was assessed by SDS-PAGE followed by altered Coomassie staining (10% acetic acid only). (**B**) Samples for electron microscopy were prepared the same as in A, with Hsp12 at 5 ug/ml. Images where taken prior to centrifugation. Samples where spread on glow-discharged EM grids and stained using 2% uranyl acetate.

### Desiccation protection is not conserved among all hydrophilins

To address whether the remarkable features of Hsp12 were common to other yeast hydrophilins, we examined a second yeast hydrophilin, Stf2 (Garay-Arroyo et al., 2010). Expressing high levels of Stf2 in exponentially dividing cells did not significantly improve their desiccation tolerance and failed to provide any synergistic tolerance with trehalose (Figure 4A). Additionally, deletion of Stf2 alone (*stf2∆*), or in combination with a loss of trehalose synthesis (*tps1∆ stf2∆*) had no effect on the short-term desiccation tolerance of stationary phase yeast, unlike the pronounced increase in desiccation sensitivity displayed by *tps1∆ hsp12∆* cells (Figure 1A and 4A). Unlike Hsp12, Stf2 also did not exhibit any detectable secondary structure in the presence or absence of DMPG (Figure 4B). Additionally, the vesiculation activity of Stf2 varied greatly from what we observe with Hsp12. While Hsp12 clearly demonstrates membrane vesiculation activity, Stf2 seems to lead to possible vesicle aggregation, even at low concentrations (Figure 4C-D). These functional differences between Hsp12 and Stf2 suggest that hydrophilins likely carry out distinct biological and molecular functions, despite sharing the general physical properties. Moreover, the biological and biochemical differences between Sft2 and Hsp12 further support the view that the specific membrane remodeling activity of Hsp12 contributes to its desiccation tolerance-promoting function.

**Figure 4.**
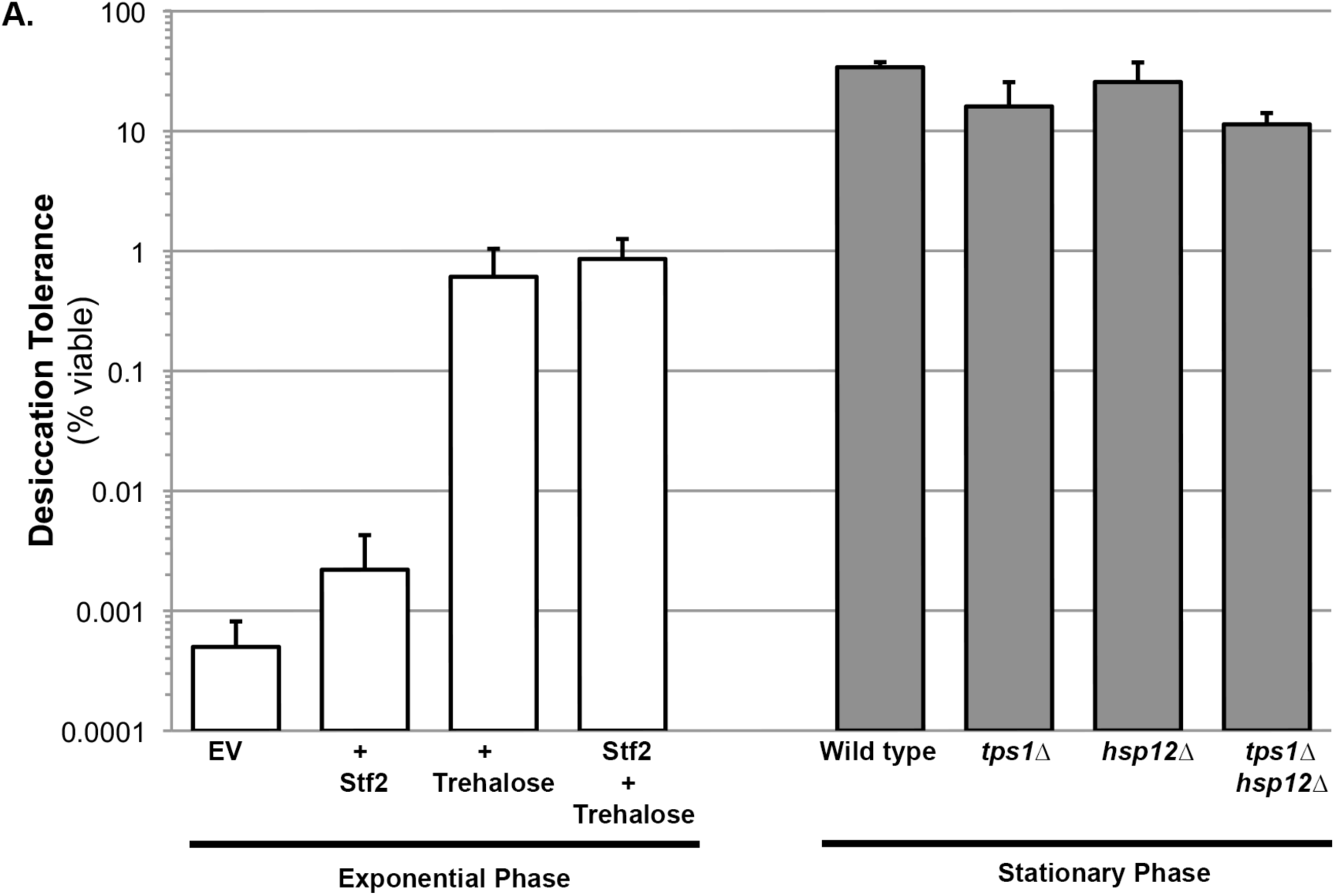
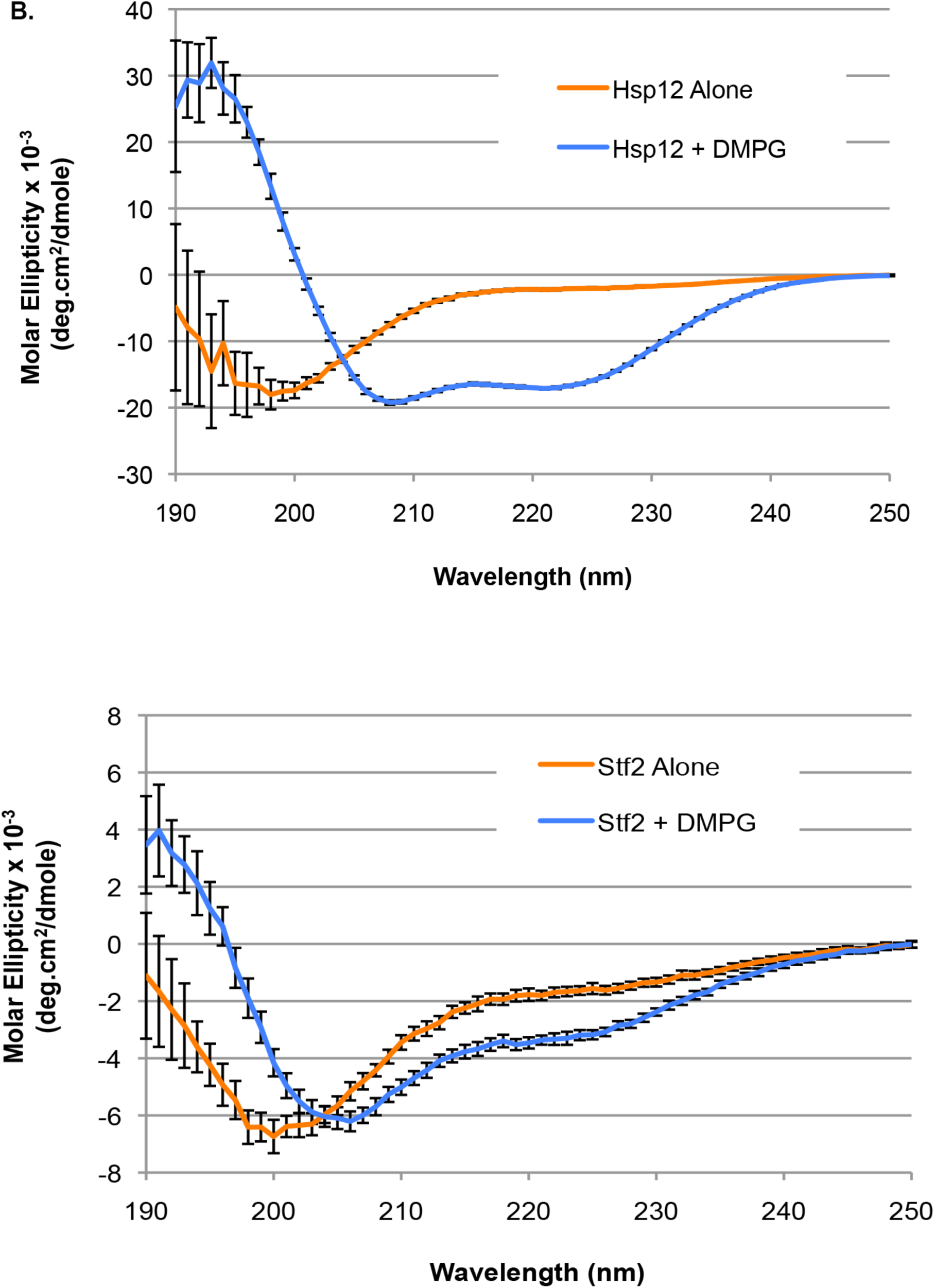
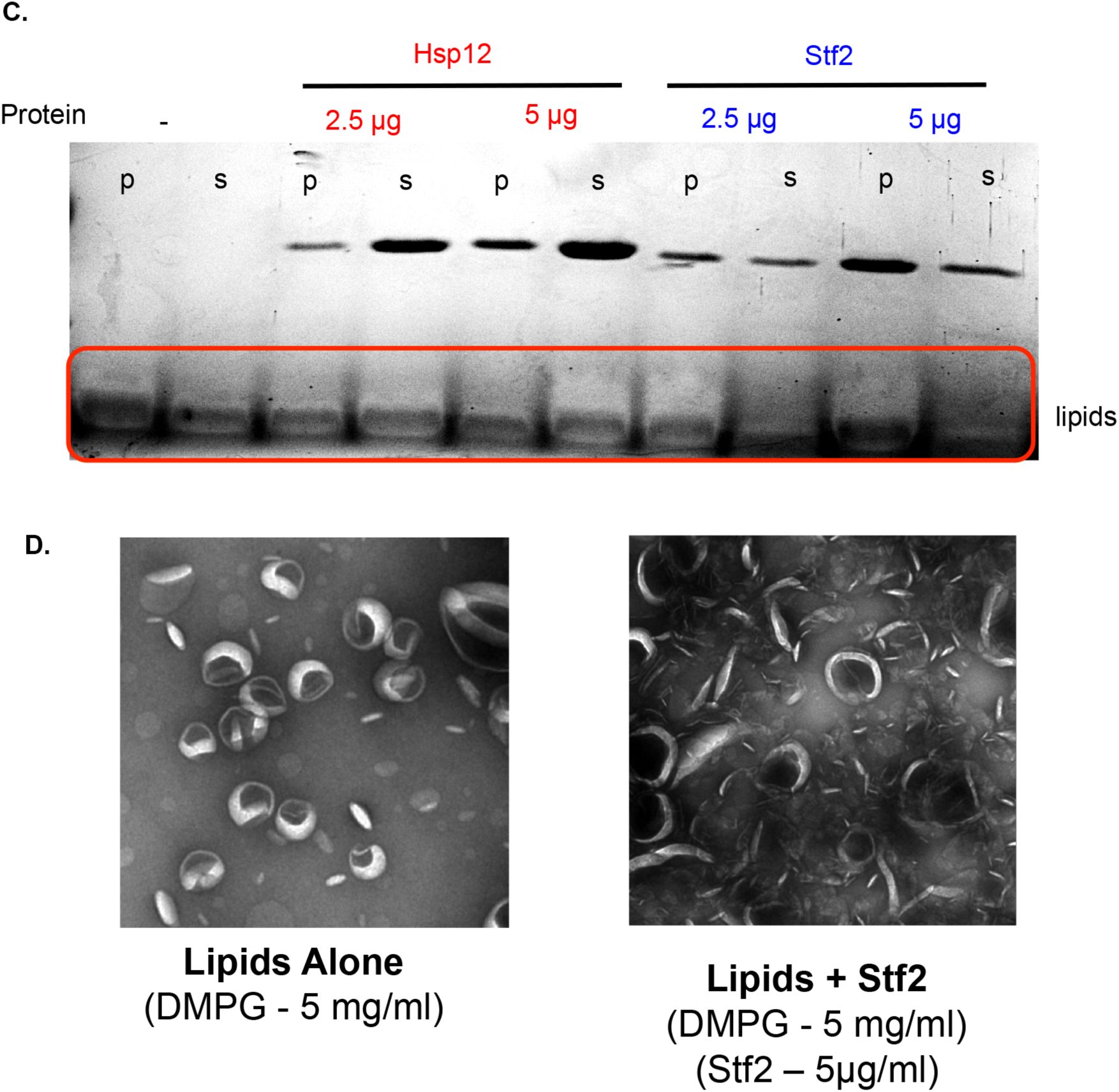
Desiccation protection not a common hydrophilin feature. (**A**) <u>Exponential Phase</u>. Yeast cells were grown to mid-exponential phase (OD < 0.5) in 728 selective media (SC-Ura+Dextrose). Cells were then transferred to selective media with trehalose (SC-Ura+Dextrose, 2% trehalose) for 1 h. Cells were collected, washed, and air dried for 2 d at 23 °C, 60% relative humidity (RH), then rehydrated and assessed for viability 731 by counting colony forming units (CFU). Yeast cells used for exponential-phase experiments performed in strain HTY163 (*nth1∆* (trehalase delete)), additionally: + EV (HTP82 - p426-*TDH3pr)*, + Stf2 (HTP163*), +* Trehalose Transporter (*TDH3pr-AGT1*), or both Stf2 and Trehalose Transporter (HTP163 and *TDH3pr-AGT1*). <u>Stationary Phase</u>. Yeast cells were grown to saturation (5 days), air-dried for 2 days at 23 °C, 60% relative humidity (RH), then rehydrated and assessed for viability by counting colony forming units (CFU). Desiccation tolerance of wild type (BY4742), HTY104 (*tps1∆*), HTY14 (*stf2∆*) or HTY179 (*tps1∆stf2∆*) cells (**B**) Circular dichroism spectroscopy performed on 0.32 mg/ml Hsp12 or 0.32 mg/ml of Stf2 in the presence or absence of 1.2 mg/ml DMPG small unilamellar vesicles (SUVs), measuring from 250 to 190 nm. Units converted to Mean Molar Residue Ellipticity, accounting for concentration and protein size. (**C**) DMPG liposomes (5 mg/ml) where incubated for 1 hour at room temperature in the presence or absence of Hsp12 (2.5-5 µg/ml) or in the presence of Stf2 (2.5-5 µg/ml). Pellet (p) and supernatant (s) fractions where separated by high-speed centrifugation, lipid distribution was assessed by SDS-PAGE followed by altered Coomassie staining (10% acetic acid only, deeper stain used to visualize lipids in cleared stain). (**D**) Samples for electron microscopy where the same as in A, with Stf2 at 5 µg/ml. Images where taken prior to centrifugation. Samples where spread on glow-discharged EM grids and stained using 2% uranyl acetate.

## DISCUSSION

We exploited the conditional desiccation tolerance of yeast to provide important new insights into the stress effectors of desiccation tolerance. First, we demonstrated a dramatic synthetic loss in desiccation tolerance in stationary cells that were deleted for *HSP12*, a hydrophilin, *and TPS1*, a biosynthetic enzyme of trehalose. This synthetic sensitivity reveals a function for this hydrophilin in desiccation tolerance that was missed because it was masked by an overlapping function with trehalose. The function of Hsp12 as a stress effector of desiccation was supported further by its ability, when highly expressed, to confer partial desiccation tolerance to exponentially dividing cells. While Hsp12 has previously been reported to mitigate lethality due to heat stress and osmolarity, we were unable to repeat those findings (Figure S1) (Welker et al., 2010). Furthermore, the previous study used a temperature for heat shock of 58°C, a temperature that budding yeast is unlikely to experience neither in the wild nor fermenting vats (Welker et al., 2010). In contrast, desiccation occurs readily in nature, so the desiccation tolerance conferred by Hsp12 is likely one of its true physiological functions. These results, coupled with our previous demonstration for the importance of a subset of tardigrade hydrophilins in desiccation tolerance, provide compelling evidence for the causal roles of a hydrophilin in desiccation tolerance in most anhydrobiotes.

Hsp12’s lipid-induced folding, its *in vitro* membrane remodeling activity, and its *in vivo* role in desiccation tolerance are not shared with Stf2, another yeast hydrophilin. These features of Hsp12 provide new insights into hydrophilins and stress biology. Like yeast, most organisms have large families of hydrophilin-like genes. The distinct causal role of Hsp12 in desiccation tolerance reveals that the generic properties of hydrophilins of charge, size and intrinsic disorder, are not sufficient to mitigate the stresses of desiccation. Thus, the other hydrophilins like Stf2 must have other unrelated functions. One simple idea is that these other functions mitigate yet to be determined stresses. If so, hydrophilins may be a major new class of diverse stress effectors.

A second major insight from this study is the demonstration that trehalose and Hsp12 can synergize to counteract all the major stresses that yeast cells encounter upon severe water loss. Exponentially dividing cells increased their desiccation tolerance 1000-fold when they were engineered to have levels of Hsp12 or trehalose similar to levels in stationary cells. This dramatic but partial desiccation tolerance was increased another 60-fold in exponentially dividing cells expressing both trehalose and Hsp12, surpassing the tolerance seen in stationary phase cells. The fact that this simple sugar and small protein together can confer complete tolerance to sensitive cells speaks to their potency to mitigate major cellular stresses. Importantly, given their small size and relatively simple structure, the molecular mechanisms underlying their stress-effector activities are likely much simpler than the more conventional, structurally and functionally complex stress effectors. The inferred simplicity of their molecular activities may be important in conditions of desiccation where water and energy dependent activities are limited.

However, we also show that compared to the *hsp12∆* or *tps1∆* single mutants, the *tps1∆ hsp12∆* double mutant was synergistically impaired in its ability to propagate a membrane prion [*GAR*^+^] in dividing aqueous cells. This result suggests that the molecular activities of these two factors in membrane/membrane-protein stasis are not confined to the desiccation stress. Rather, they can act together to modulate membrane or membrane-protein function in aqueous conditions, perhaps providing a new clue for the reason why their regulation is linked in response to numerous stresses.

We provide *in vivo* and *in vitro* evidence that both trehalose and Hsp12 can modulate proteostasis in response to desiccation. We show that they prevent desiccation-induced aggregation of luciferase *in vivo* and of citrate synthase *in vitro*. Trehalose and Hsp12 also synergize, albeit weakly, to protect luciferase activity *in vivo*, and individually they protect citrate synthase activity *in vitro*. Whether the stabilization of enzyme activity is different or just another manifestation of their anti-aggregation activity is not clear. Interestingly, the proteostasis functions of trehalose and Hsp12 are not identical as cells lacking only trehalose are compromised for [*PSI*^+^] and [*GAR*^+^] propagation while lacking Hsp12 are not. These differences likely indicate that trehalose and Hsp12 may modulate proteostasis by different mechanisms.

In addition to Hsp12’s proteostasis activities, we show that it has an *in vitro* activity of membrane remodeling as evidenced by its vesiculation of lipid vesicles. This membrane activity is consistent with several *in vivo* and *in vitro* observations. Hsp12 synergizes with trehalose to allow propagation of the membrane prion [*GAR*^+^] and also acquires secondary structure upon binding to lipids. A previous study suggested that cells lacking Hsp12 have altered membrane structure and fluidity (Welker et al., 2010).

Interestingly, trehalose prevents abnormal membrane vesiculation in desiccated nematodes (Erkut et al., 2011). *In vitro* experiments have suggested that trehalose intercalates into membranes to alter their melting properties (Crowe et al., 1984; 1987; Leslie et al., 2002; Tang et al., 2007). Based upon these observations and our observations of Hsp12 in this study, we propose that, alongside protecting against proteotoxicity, Hsp12 and trehalose cooperate in membrane homeostasis. We propose that desiccation induces membrane damage. This damage is prevented by the intercalation of trehalose into membranes. If damage occurs, it is removed by Hsp12’s membrane remodeling activity. These distinct activities explain the synergistic requirement for trehalose and Hsp12 in both short-and long-term desiccation tolerance. Additional genetic and biochemical experiment will be needed to test this model.

Regardless of their mode of function, the fact that just trehalose and Hsp12 are sufficient to mitigate the major stresses of desiccation has major implications for engineering desiccation/drought tolerance in other organisms. Water performs so many important functions in biology; it was easy to imagine that the removal of water would generate many stresses that require many stress effectors to combat. The need for a plethora of stress effectors would make transmitting this trait to another organism too complex. The ability to generate desiccation tolerance with only two factors makes this engineering eminently more feasible. Attempts to engineer plants to synthesize more trehalose have been met with technical difficulties because of the sugar's additional roles in metabolism. Our results provide new impetus to overcome those hurdles. Furthermore, Hsp12 has no known metabolism side effects. Given the sufficiency of Hsp12 alone to confer partial desiccation tolerance, it will be very interesting to test whether Hsp12 by itself might confer drought tolerance in plants.

## MATERIALS AND METHODS

### Strains and Growth Conditions

Standard yeast propagation and transformation procedures were used. Yeast strains are described in Supplemental Table 2. Strains were grown in nonselective (YP, 1% yeast extract and 2% peptone) or selective (synthetic complete, SC) media containing 2% glucose. Cultures were grown to saturation from a single colony by incubating cultures 5 days at 30°C for stationary phase experiments. Cultures where grown to an OD – 0.5 for mid-log experiments. All experiments were repeated at least two times on separate days with separate isolates when appropriate. Hsp12 and Stf2 over-expression was driven from a plasmid under a constitutively expressed promoter (p423-*TDH3pr*-*HSP12* (HTP126) or p426-*TDH3pr-STF2* (HTP163)) or carry an empty vector plasmid (p423-*TDH3pr* (HTP64), p426-*TDH3pr* (HTP82)).

### Desiccation Tolerance Assay

Saturated Cultures: Approximately 10^7^ cells were withdrawn from liquid cultures and washed twice in dilute water and then brought to a final volume of 1 ml. Undesiccated controls were plated for colony counting. Two hundred microliter aliquots were then transferred to a 96-well tissue culture plate (Becton Dickinson, 353075) and centrifugated, and water was removed without disturbing the cell pellet. Cells were allowed to desiccate in a 23°C incubator with a constant 60% relative humidity (RH), with the lid raised, for at least 48 hr. Long-term desiccation experiments were kept for indicated time periods in a 96-well tissue culture plates at 23°C, 60% RH. Samples were resuspended in assay buffer and plated for colony counting.

Logarithmic Samples: Cells were grown to mid-exponential phase (OD < 0.5) in different selective media, depending on strain and plasmid necessities. Cells were then transferred to inducible media: ± 2% trehalose, for 1 h. Following induction, ∼10^7^ cells were withdrawn from liquid cultures and same parameters for drying where used as with saturated cultures. Data were entered into a spreadsheet (Microsoft Excel 2008 for Mac version 12.3), and the number of colony forming units per milliliter (cfu/mL) for each plate was computed. For each experiment, number of colony forming units per milliliter for the two controls was averaged. The relative viability of each of the two experimental samples was determined by dividing the number of colony forming units per milliliter for that sample by the average number of colony forming units per milliliter of the undesiccated control. These two relative viability values were then averaged and their SD was computed using the STDEVP worksheet function.

### Luciferase Assay

Yeast cells bearing 2µ plasmids that direct the expression of a temperature-sensitive firefly luciferase protein from the constitutive glyceraldehyde-3-phosphate (*TDH3pr*) promoter (HTP111 - p426-*TDH3pr*-FFL) or empty vector (HTP82 - p426-*TDH3pr*), were grown to mid-log phase in SC-Ura-His. Luciferase activity was measured *in vivo* by addition of 0.5 mM D-Luciferin (Sigma) to equal number of intact cells. Light emission was measured immediately with a TD-20/20 Luminometer (Turner Designs, Sunnyvale CA). Desiccated samples were air dried for two days followed by rehydration in SC-His - Ura + cycloheximide (10 µl/ml, to block new FFL protein synthesis) and subjected to the same assay treatment after rehydration. Measurements are reported as Relative Light Units. Using glass bead lysis, total cellular protein was extracted. After removing unbroken cells by low-speed centrifugation (1,000g for 3 min), cleared lysates were spun at high speed (350,000g for 10 min, Beckman Coulter TLA-100) to collect insoluble aggregates. Total protein from cleared lysates and high-speed supernatants were then followed by SDS-PAGE and reacted with antiserum recognizing luciferase (Promega). All experiments were repeated three times on separate days with separate isolates.

### Citrate Synthase Assay

Citrate Synthase from Porcine Heart ammonium sulfate suspension (C3260) was purchased from Sigma Aldrich. Suspension was centrifugated at 14,000 rpm for 10 minutes at 4°C. Supernatant was discarded and the pellet was resuspended in 750 µl MilliQ water. To remove residual ammonium sulfate, citrate synthase was desalted in MilliQ water using a 5 ml HiTrap Desalting Column (GE Healthcare). Fractions were evaluated for protein content via staining 30 µl of each fraction with 100 µl of Coomassie protein assay reagent (Sigma) diluted 1:5 in MilliQ water. Fractions with detectable protein content were pooled and concentrated using an Amicon Ultra 0.5 ml 10K centrifugal filter unit (Merck Millipore Ltd). Final concentration determined via measuring absorbance at 280 nm on a Nanodrop 2000 spectrophotometer (Thermo Scientific).

Citrate synthase aggregation and enzymatic activity assay:

Citrate synthase (CS) and protectants were added to MilliQ water to achieve final concentration of 1.2 mg/ml CS, and varying concentrations of protectants in 100 µl. For measuring enzymatic activity, 1 µl of each sample was diluted by a factor of 17 in MilliQ water. 2 µl of this dilution was then mixed with 198 µl of Enzymatic Activity Assay Solution [100mM oxaloacetic acid (Sigma-Aldrich), 100mM 5,5’-Dithiobis(2-nitrobenzoic acid) (Sigma-Aldrich), 150mM Acetyl Coenzyme A (Sigma-Aldrich)] and absorbance at 412 nm was measured immediately and after 120 seconds in a 50 µl Micro Cell cuvette (Beckman Coulter) with measurements made with a 720 UV/Vis spectrophotomer (Beckman Coulter) equipped with a 50 µl Micro Cell adapter (Beckman Coulter). After measurement, samples were moved to 1.5 ml microcentrifuge tubes and set to dry in a Centrivap Concentrator (Labconco) set to 35°C and 29 bar for 24 hours. Samples were then moved to a 23°C incubator and allowed to incubate open to atmospheric pressure for an additional 5 hours. Samples were rehydrated in 100 µl MilliQ water, resuspended vigorously 5 minutes after addition of water. 2 µl of this resuspension was added to 198 µl Enzymatic Activity Assay Solution, mixed and moved to a cuvette, measuring absorbance at 412 nm immediately and after two minutes. Change in absorbance across two minutes after drying compared to before drying to achieve relative activity. For measuring aggregation, absorbance at 340 nm was measured for each sample before and after drying. Drying and rehydration performed as described for enzymatic activity assay. Measurements after drying subtracted from before drying measurements to achieve change in aggregation after drying. Cuvette was washed with MilliQ water and 95% ethanol between all measurements.

### Expression and purification of Hsp12p

BL21 *E. coli* cells transformed with a pET-15B vector with HSP12 behind a galactose-inducible promoter grown overnight in 30 ml cultures in LB with 100 ng/ml ampicillin. Transformants were grown up overnight in 30 mL cultures in LB with 100 ng/ml ampicillin at 37° C. 20 mL of this culture were added to 2L of LB with ampicillin and incubated at 37° C until culture reached an OD600 of 0.5-0.9, as determined by spectrophotometry using a 720 UV/Vis spectrophotomer (Beckman Coulter). HSP12 expression induced by addition of IPTG (Isopropyl β-D-1-thiogalactopyranoside) to final concentration of 1 mM and incubation for an additional 4 hours at 37°C. Cultures were then pelleted by centrifugation at 4,000 rpm for 30 minutes at 4°C. Discarded supernatant and resuspended cells in 50 mL of 1x PBS. Cells were frozen with liquid nitrogen, and stored at −80°C until use. Pellets were isolated and resuspended in lysis buffer (2mM EDTA, 2mM DTT, 20mM HEPES, pH 7.4) then sonicated using a blunt tip sonicator set to 30% amplitude, 20 minutes, pulse 1 second on, 1 second off at 4° C. Sonication-lysed cells were then centrifugated and supernatant was collected. This sample was then boiled at 95° C for 15 mins and immediately moved to ice. A pale yellow precipitate was separated away from the proteins that are still soluble after boiling by centrifugation. Supernatant was then applied to a HiTrap Capto-Q ImpRes anion exchange column (GE Healthcare Life Sciences), and washed with lysis buffer with increasing concentrations of NaCl. Hsp12 was eventually eluted by buffer between 7.5 mM and 10 mM NaCl. The protein was then concentrated using a MW3000 filtration unit, aliquoted, frozen in liquid nitrogen, and stored at −80° C until use (Supple Fig. 2).

### Expression and purification of Stf2p

STF2 fused to GST was cloned into a pGEX6 vector behind a galactose-inducible promoter and transformed into BL21 cells. Transformants were grown overnight at 37° C in 30 mL of LB liquid media, selecting for plasmid with 75 ng/ml ampicillin. 2 L LB cultures with 75 ng/ml ampicillin were inoculated with 20ml of these overnight cultures and set to 37° C and 140 rpm in an Innova 4330 incubator (New Brunswick Scientific). After 4 hours, expression of STF2 was induced with 4 mL of 0.1M IPTG and cultures were allowed to incubate overnight at 18° C. Cells were harvested and lysed in lysis buffer 2 (150mM NaCl, 2mM EDTA, 2mM DTT, 20mM HEPES, pH 7.4) using the same sonication method as above (see Purification of Hsp12). Then, lysed cells were centrifugated and the supernatant was collected. 1.5 ml of Pierce Glutathione Agarose beads from Thermo Fisher Scientific were added and incubated in the supernatant for an hour at 4° C. Beads were washed in 3 washes in lysis buffer 2, 3 washes in lysis buffer 3 (500mM NaCl, 2mM EDTA, 2mM DTT, 20mM HEPES, pH 7.4), and 3 washes in lysis buffer 2, and resuspended in lysis buffer 2. Prescission protease was added to washed beads and this mixture was incubated at 4° C overnight. The mixture was then centrifugated and the supernatant was collected, concentrated with an Amicon Ultra-4 Ultracel 3K centrifugal filter unit from Millipore, aliquoted, frozen in liquid nitrogen, and stored at −80° C (Supple Fig. 2).

### Prion phenotypic assays

To asses prion state, cells were plated on media lacking adenine (SC-ADE) for [*PSI+*] strains, or on YP media with 0.05% Glucosamine and 2% Glycerol for [*GAR+*] growth was compared to their ability to grow on rich YP-GAL media. To determine % Prion = 100 X CFU Desiccated Cells (Prion Selective Media) / CFU Desiccated Cells (Non-Selective Rich Media).

### Liposome preparation and vesiculation

DMPG liposomes (Avanti Polar Lipids (840445) in 100 mM NaCl, 20 mM Tris-HCl pH 7.4 were sonicated for 2 minutes in a water bath at room temperature. Liposomes were incubated with protein for 60 minutes and centrifugated at 250,000g for 15 min in a Beckman Coulter TLA-100 rotor. Resuspended pellets and supernatants were analyzed by SDS-PAGE. Gels were stained with 0.1% Coomassie in 10% Acetic acid and destained in water. For EM, samples were spread prior to centrifugation on glow-discharged electron microscopy grids and stained using 2% uranyl acetate and viewed in a Joel 1200 EX Transmission Electron Microscope.

### Preparing liposomes for circular dichroism

1,2-dimyristoyl-*sn*-glycero-3-phospho-(1'-rac-glycerol) (DMPG) from Avanti Polar was stored in chloroform at −80° C in glass tubes until used. DMPG aliquots were thawed and aspirated using a stream of nitrogen gas to form a clear, dry film on the inside surface of the tube. Tubes were then placed inside a vacuum desiccator and sealed. Vacuum was then applied to the desiccator, and the samples were left at room temperature under vacuum overnight to remove any remaining chloroform. Then, they were resuspended in 10mM sodium phosphate, pH 7.4 and moved to a 1.5 ml plastic microcentrifuge tube. Resuspended aliquots were then vortexed until a homogenous, cloudy white solution was formed. Tubes were placed on ice in a 4° C room, and sonicated using a probe tip sonicator equipped with a blunt tip and set to 20% amplification, 7.5 minutes, with pulses of 2 seconds on and 2 seconds off. Lipids were then spun down at 14,000 rpm at 4° C for 10 minutes to precipitate any metal shards from sonication. Supernatant was then moved to a new 1.5 ml microcentrifuge tube and left on ice until use in circular dichroism.

### Circular dichroism spectra

Circular Dichroism (CD) spectra were obtained using an Aviv 2000 Circular Dichroism Spectrometer Model 410 (Aviv Biomedical Inc.). CD signal was measured from 250 to 190 nm at 25° C, averaging measurements at every nm for 10 seconds. Hydrophilins and DMPG were kept on ice until added to samples. Once added, samples were mixed well by flicking and incubated at room temperature for 10 minutes before spectra were measured. A 1M stock of trehalose and 100mM stock of SDS were prepared in 10mM sodium phosphate, pH 7.4 and added to achieve desired concentrations. Proteins were added to samples to achieve a concentration of 0.32 mg/ml. For samples with DMPG, DMPG was added to achieve a concentration of 1.32 mM (1.2 mg/ml. Samples were made at a volume of 400 µL and 300 µL were pipetted into a 0.1 cm cuvette for CD measurements. The cuvette was washed with water and ethanol between measurements, drying completely each time. Samples with protein were blanked against those with the same buffer and concentration of DMPG without hydrophilins. All spectra were converted to units of mean residue molar ellipticity before plotting on graph.

### SGA Strain Construction

Strains were construction according to published SGA protocol from Tong et al. with minor alterations (Tong et al., 2001). In brief, *tps1∆* was grown overnight as well as every strain in the ATCC deletion collection. Using a 48-density prong, *tps1∆* was pinned onto solid YEP + Galactose media and the deletion collection were pinned on top. This was allowed to grow overnight at 30°C. Cells were then replica plated onto YEP + Galactose + G418 (100 mg/mL) + Hygromycin (100 mg/mL) to select for diploids. These were grown at 30°C overnight. Diploids were then replica plated onto sporulation media (10g Potassium Acetate, 1g Yeast Extract, 0.5g Galactose, 0.1g amino acid Histidine, Lysine, Leucine, and Uracil supplement, Zinc Acetate, 0.2g Raffinose). Cells were sporulated for 5 days. After sporulation, we selected for MATa haploids on SC + Galactose – Histidine – Arginine + Canavanine (50 mg/L) + Thialysine (50mg/L). Then we selected for G418 resistance on SC + Galactose – Histidine – Arginine + Canavanine (50 mg/L) + Thialysine (50 mg/L) + G418 (100 mg/mL). Finally, we selected for Hygromycin resistance on SC + Galactose – Histidine – Arginine + Canavanine (50 mg/L) + Thialysine (50 mg/L) + G418 (100 mg/mL) + Hygromycin (100 mg/mL).

### High Throughput Desiccation Tolerance Assay

Single colonies of newly constructed double mutants are placed into 200 **µ**L of YEP + 2% Galactose in wells of 96-well plates and allowed to grow to saturation at 30°C with agitation. Strains were then pinned using a 48-density prong onto YEP + Galactose plates and allowed to grow for 2 days as non-desiccated controls. 20 **µ**L of each strain was also transferred into new 96-well plates to let air dry for 6 and 30 days at 23°C. Drying was done as previously mentioned. After desiccation, strains are rehydrated in 200 μL of YEP + Galactose and pinned onto solid media as before. Desiccation tolerance is assayed by comparing growth of strains after desiccation with non-desiccated controls.

### Heat Tolerance Assay

Supplemental Figure 1 (**A**) Cells (wild type, *hsp12∆*, *tps1∆*, or *tps1∆ hsp12∆)* were grown to midexponential phase (OD < 0.5) in non-selective media (YP, 1% yeast extract and 2% peptone) containing 2% glucose at 30C. Cells were plated and grown at either 30C or 37C, or grown for one hour at 34C (pre-heat shock) before growing at either 30C or 37C. (**B**) Cells (wild type, *hsp12∆*, *tps1∆*, or *tps1∆ hsp12∆)* were grown to midexponential phase (OD < 0.5) in non-selective media (YP, 1% yeast extract and 2% peptone) containing 2% glucose at 30C. Cells were then heat-shocked for 30 min at temperatures ranging from 42-60C, followed by plating and growing at 30C. Cells were also grown at 34C prior to heat shock.

## ACKNOWLEDGEMENTS

We thank Kevin A. Morano, Jeremy Thorner, Jasper Rine and Patrick Gibney for critical reading of our manuscript; members of the Koshland lab for technical support; and Reena Zalpuri at the electron microscope lab at UC Berkeley. This work was supported by a grant from the G. Harold and Leila Y. Mathers Charitable Foundation to S.X.K. and H.T., by an NIH grant (GM092813) to D.K., and by Damon Runyon Cancer Research Foundation (DRG-2137-12) to G.Ç..

## SUPPLEMENTAL DATA

**Figure S1.**
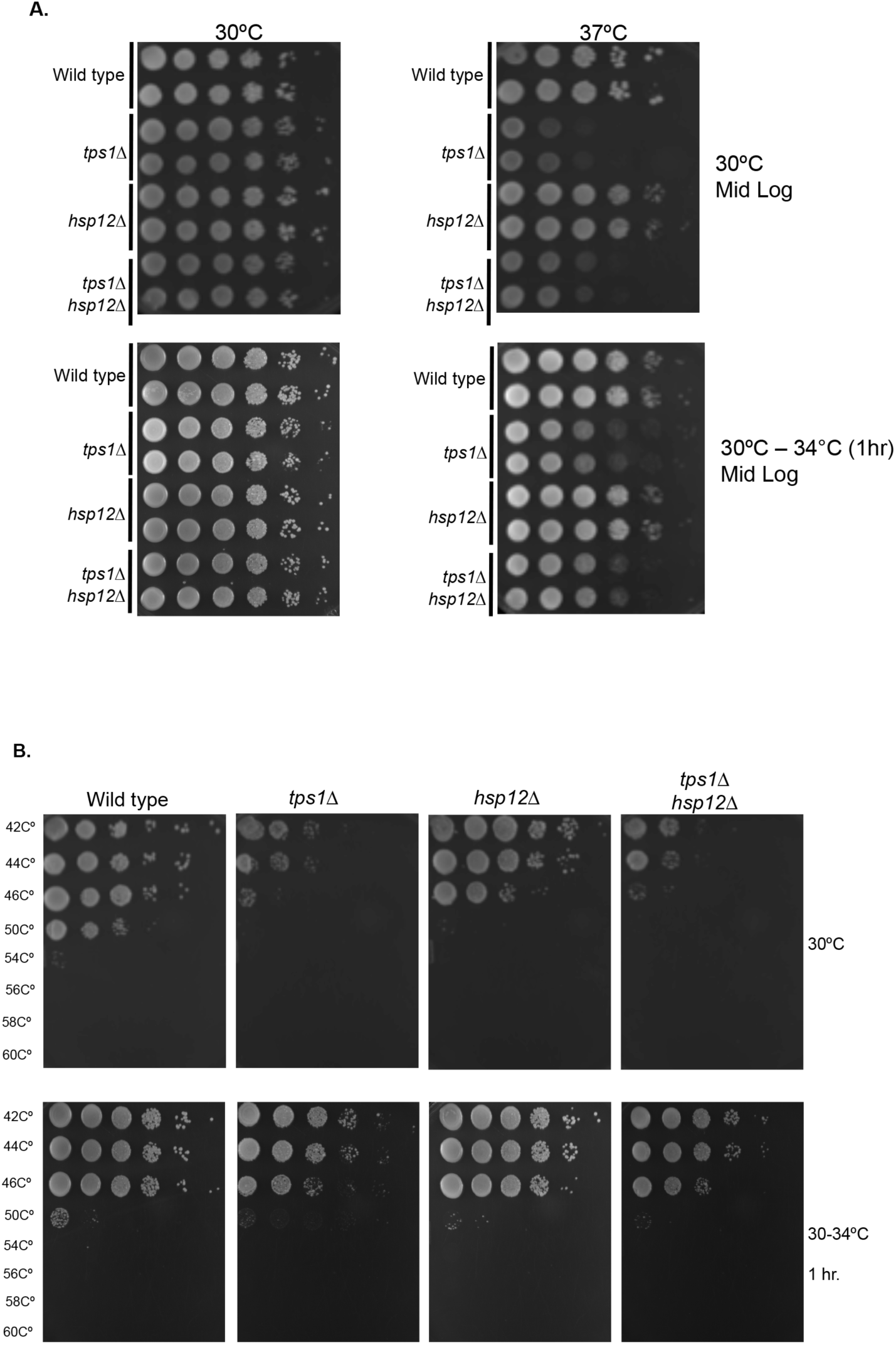
Trehalose and Hsp12 heat tolerance. (**A**) Cells (wild type, *hsp12∆*, *tps1∆*, or *tps1∆ hsp12∆*) were grown to midexponential phase (OD < 0.5) in non-selective media (YP, 1% yeast extract and 2% peptone) containing 2% glucose at 30C. Cells were plated and grown at either 30C or 37C, or grown for one hour at 34C (pre-heat shock) before growing at either 30C or 37C. (**B**) Cells (wild type, *hsp12∆*, *tps1∆*, or *tps1∆ hsp12∆*) were grown to mid-exponential phase (OD < 0.5) in non-selective media (YP, 1% yeast extract and 2% peptone) containing 2% glucose at 30C. Cells were then heat-shocked for 30 min at temperatures ranging from 42-60C, followed by plating and growing at 30C. Cells were also grown at 34C prior to heat shock.

**Figure S2.**
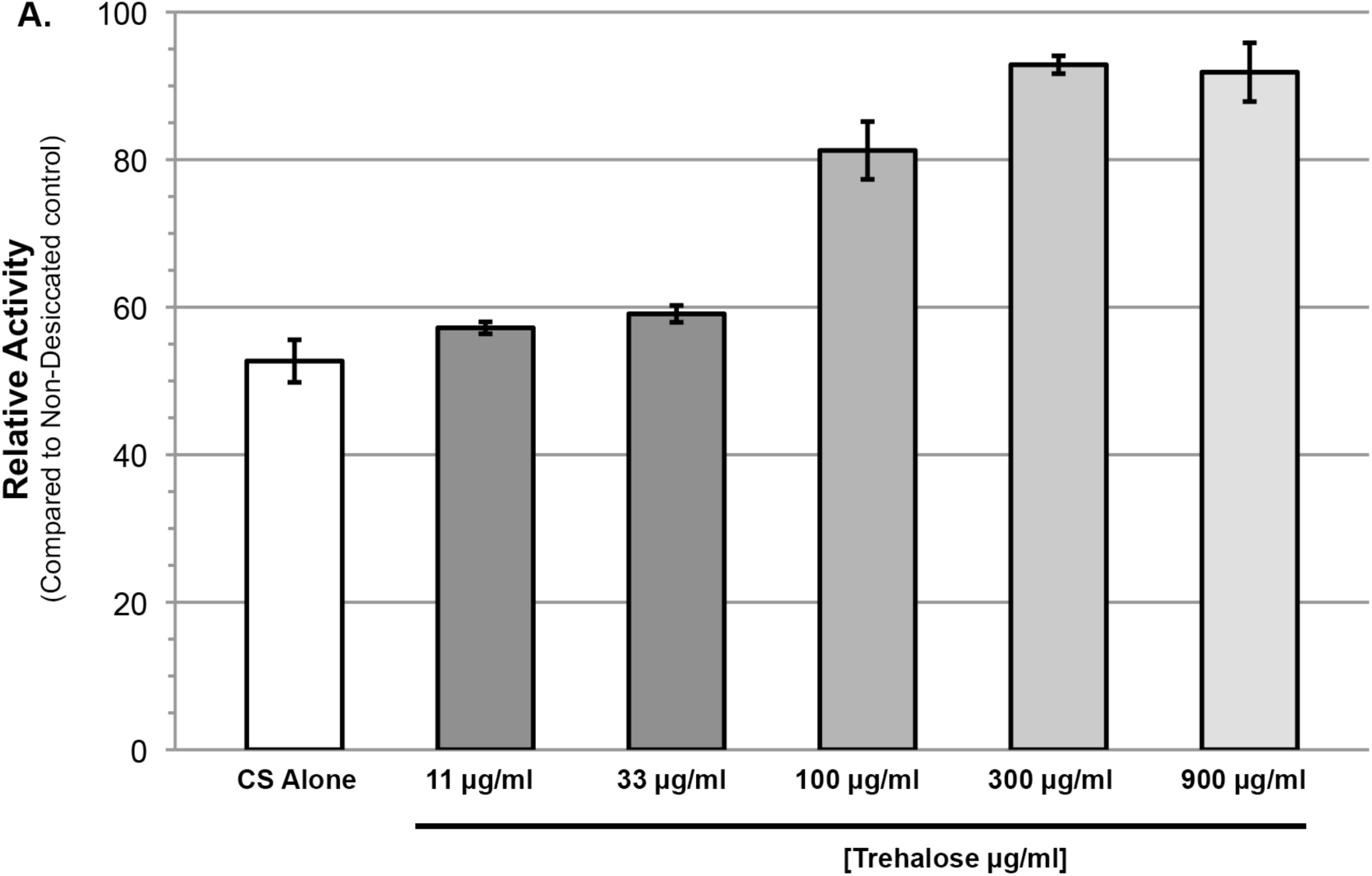
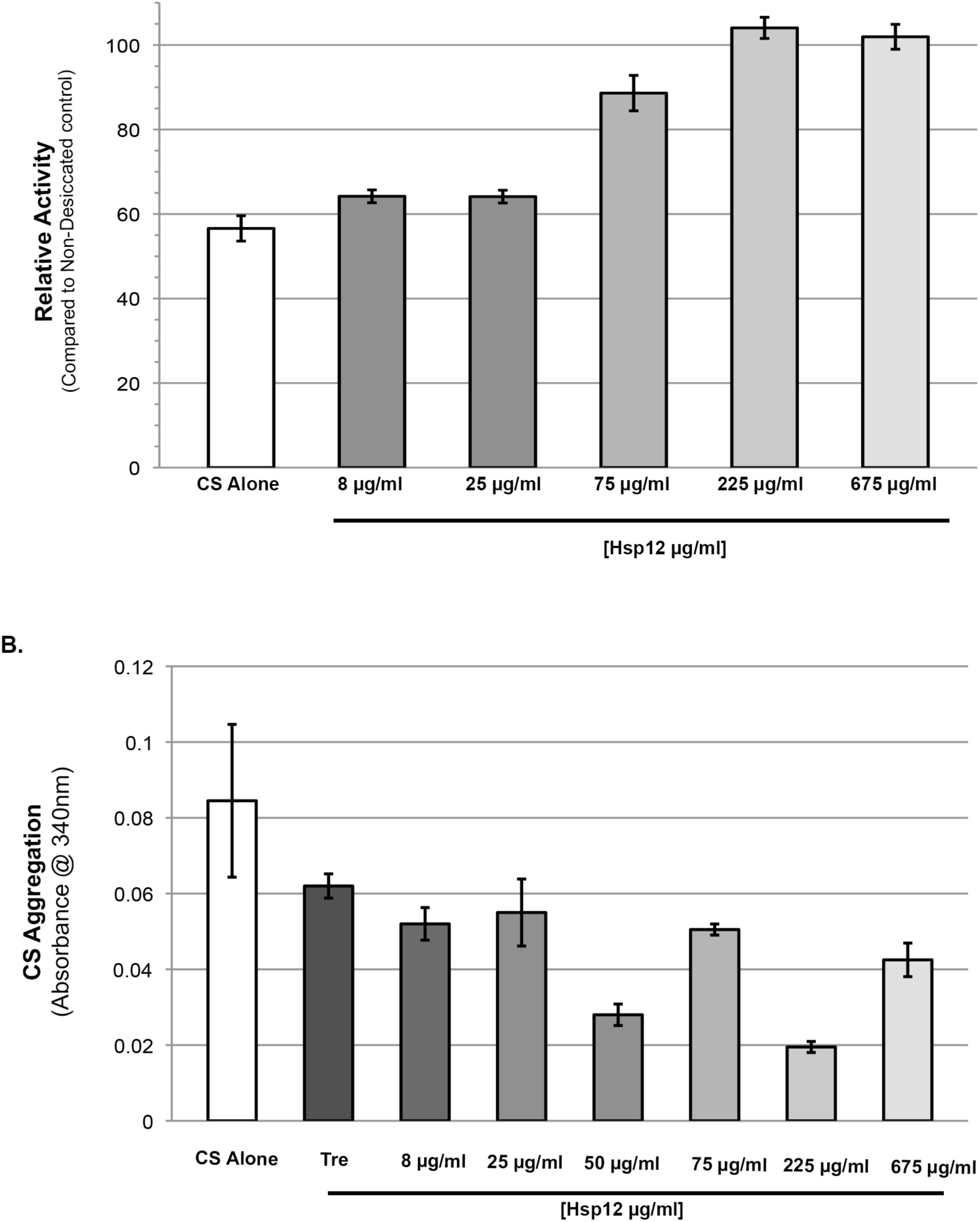
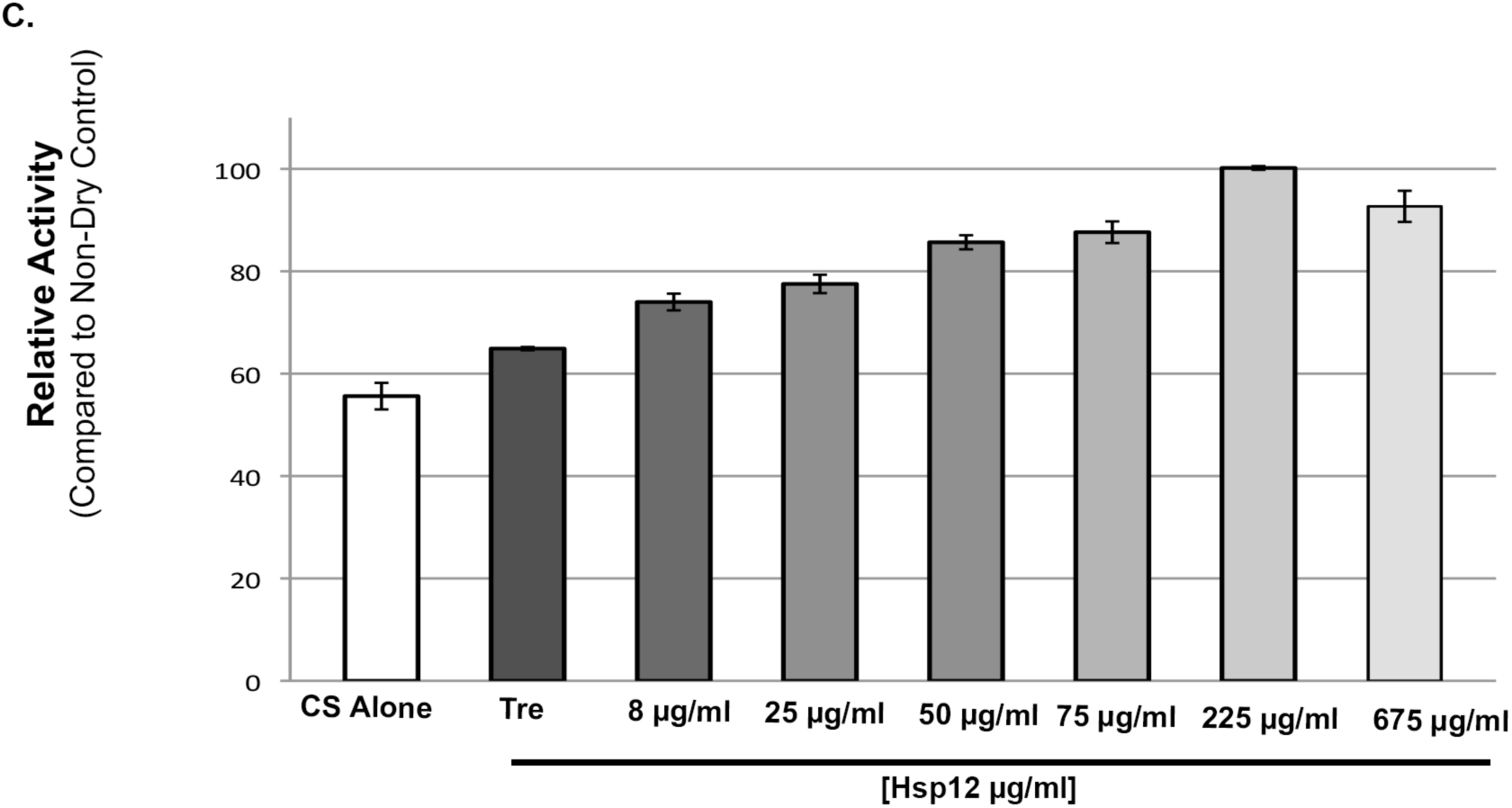
Citrate synthase activity and synergistic protection. (A) Enzymatic activity after drying of CS in presence of varying concentrations of trehalose or Hsp12. CS enzymatic was measured by examining activity-induced changes in absorbance at 412 nm after drying compared to measurements before drying. (B). Citrate synthase desiccation-induced aggregation in the presence of constant concentrations of trehalose and varying concentrations of Hsp12. Absorbance at 340 nm. Citrate Synthase (0.12 mg) and trehalose (33ug/ml). Absorbance measured after drying at 36°C for 24 hours followed by 5 hours at 23°C, then rehydration in water. Measurements after drying compared to measurements before drying. (C) Assay as in S2B, but tested for CS activity.

**Figure S3.**
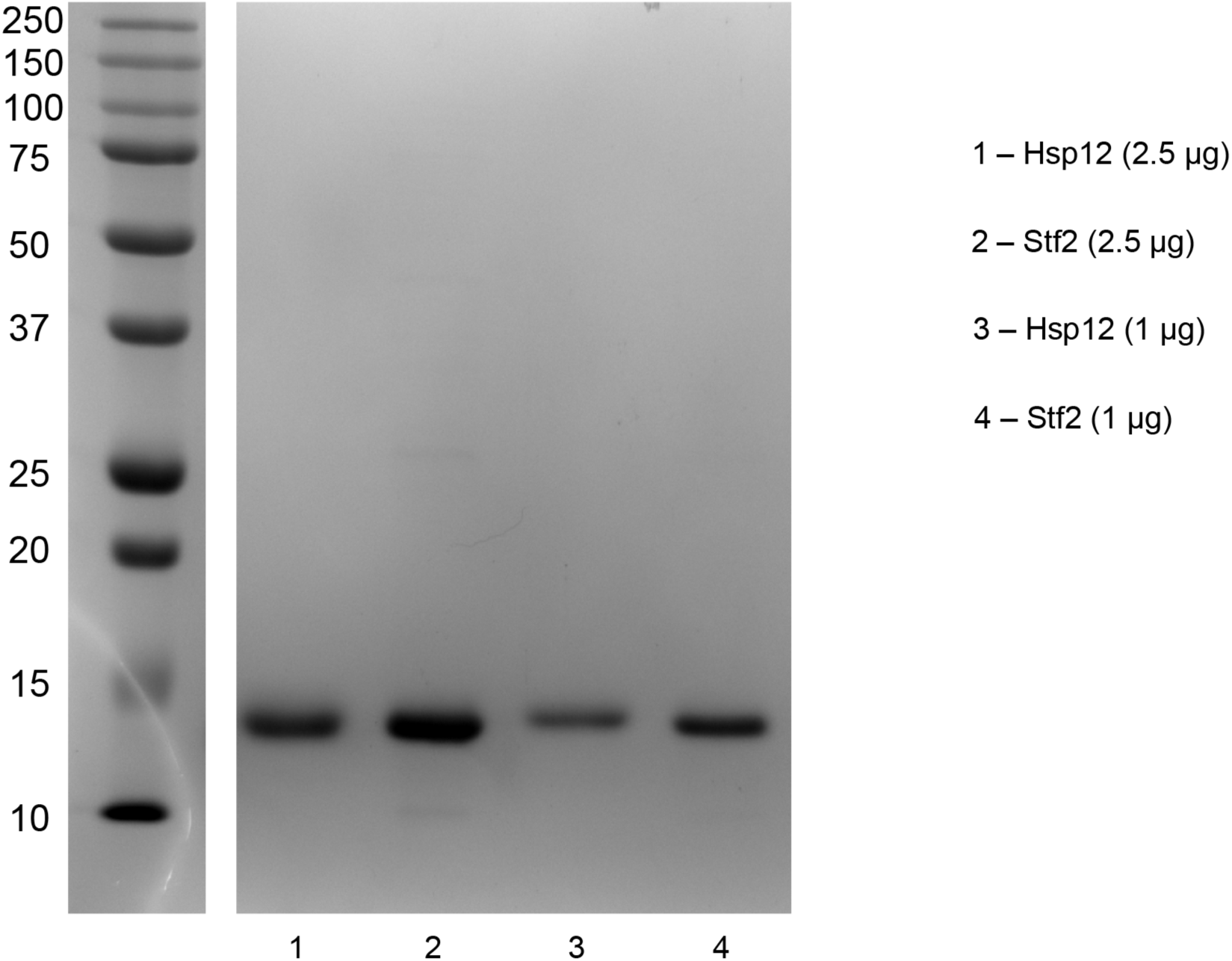
Hydrophilin protein purifaction. Hsp12 and Stf2 are bacterial derived protein preps. Gel stained with Coomassie Blue demonstrates purity of protein preps.

**Table S1.**
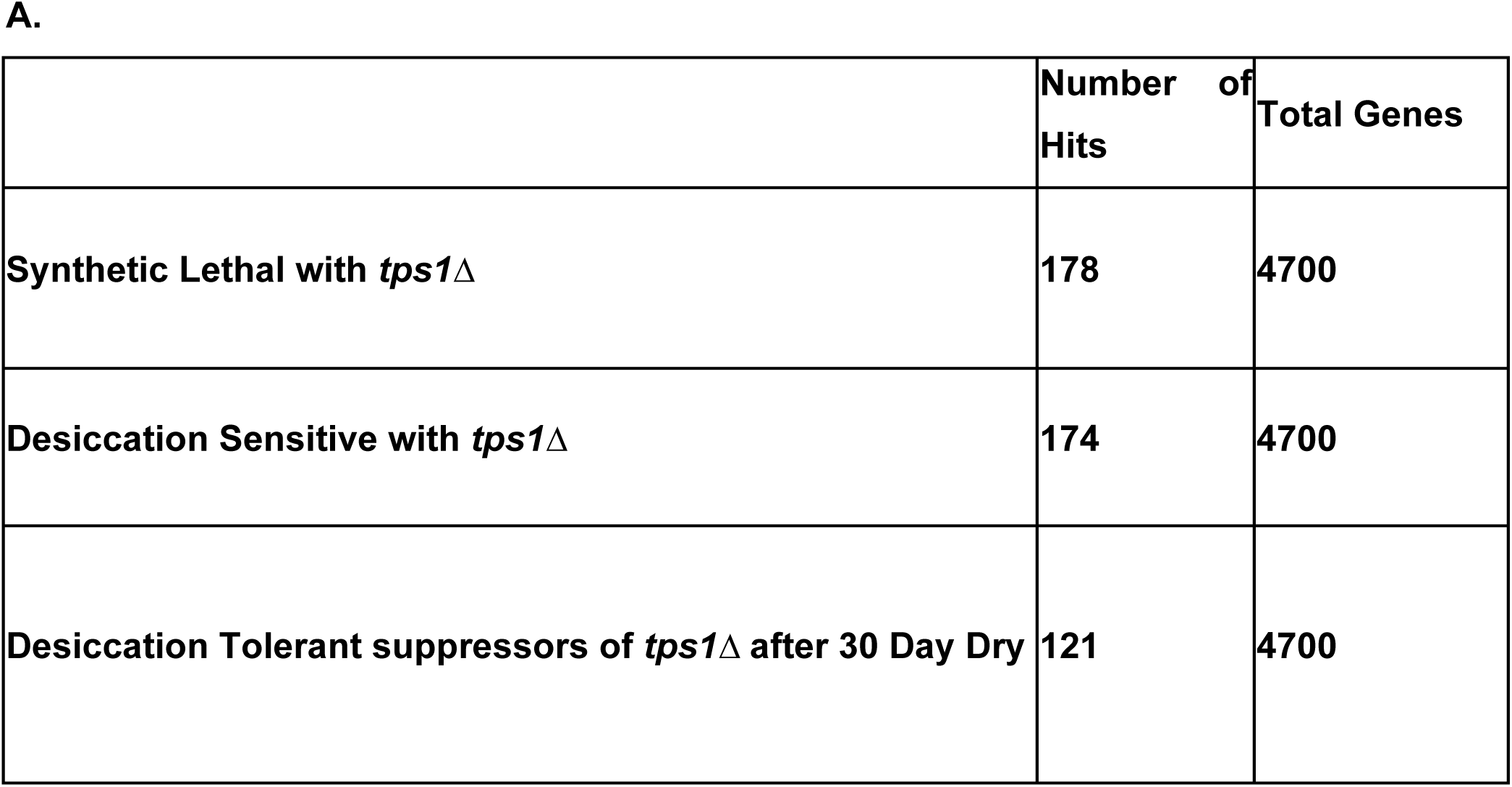

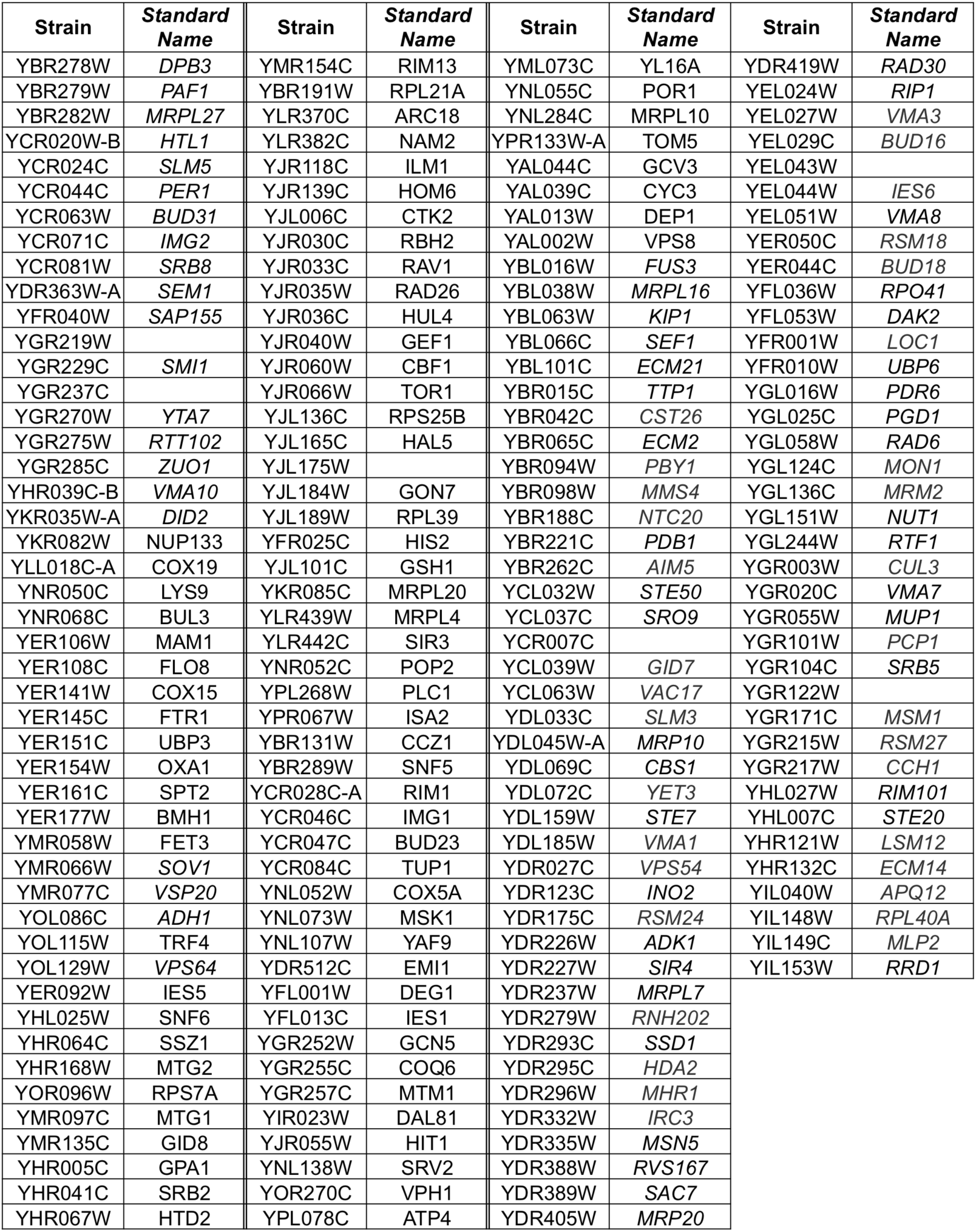
1.Synthetic Lethality with *tps1*Δ - SGA

2.Desiccation Sensitive with *tps1*Δ - 6 Day Dry - SGA

2. Desiccation Sensitive with *tps1* $\Delta$ - 6 Day Dry - SGA
| Strain | Standard Name | Strain | Standard Name | Strain | Standard Name | Strain | Standard Name |
| --- | --- | --- | --- | --- | --- | --- | --- |
| YFL014W | HSP12 | YJR052W | RAD7 | YKL137W | CMC1 | YNR002C | ATO2 |
| YBR295W | PCA1 | YJR061W |  | YKL150W | MCR1 | YNR039C | ZRG17 |
| YBR297W | MAL33 | YFL033C | RIM15 | YKL159C | RCN1 | YNR047W | FPK1 |
| YCR026C | NPP1 | YAL008W | FUN14 | YKL161C | KDX1 | YOR061W | CKA2 |
| YCR027C | RHB1 | YBL001C | ECM15 | YKL170W | MRPL38 | YOR065W | CYT1 |
| YGR256W | GND2 | YBR019C | GAL10 | YKL171W | NNK1 | YOR086C | TCB1 |
| YIR021W | MRS1 | YBR037C | SCO1 | YKL175W | ZRT3 | YOR107W | RGS2 |
| YIR023W | DAL81 | YBR203W | COS111 | YKL207W | EMC3 | YOR115C | TRS33 |
| YIR031C | DAL7 | YBR215W | HPC2 | YKR003W | OSH6 | YOR118W | RTC5 |
| YIR033W | MGA2 | YBR227C | MCX1 | YKR055W | RHO4 | YOR149C | SMP3 |
| YIR038C | GTT1 | YCR009C | RVS161 | YKR059W | TIF1 | YOR161C | PNS1 |
| YLR423C | ATG17 | YDL107W | MSS2 | YLL006W | MMM1 | YOR166C | SWT1 |
| YLR425W | TUS1/SOP10 | YDR148C | KGD2 | YLL014W | EMC6 | YOR178C | GAC1 |
| YLR447C | VMA6 | YDR194C | MSS116 | YLL019C | KNS1 | YOR182C | RPS30B |
| YER118C | SSU81 | YDR197W | CBS2 | YLL026W | HSP104 | YOR191W | ULS1 |
| YER128W | VFA1 | YDR204W | COQ4 | YLL040C | VPS13 | YOR231W | MKK1 |
| YER153C | PET122 | YDR350C | TCM10 | YLL043W | FPS1 | YOL015W | IRC10 |
| YER155C | BEM2 | YDR375C | BCS1 | YLL051C | FRE6 | YOL083W | ATG34 |
| YER167W | BCK2 | YER030W | CHZ1 | YLR025W | SNF7 | YOL084W | PHM7 |
| YMR070W | MOT3 | YFL016C | MDJ1 | YLR006C | SSK1 | YPL274W | SAM3 |
| YOL095C | HMI1 | YGL107C | RMD9 | YLR064W | PER33 | YPL273W | SAM4 |
| YOL096C | COQ3 | YGL115W | SNF4 | YLR079W | SIC1 | YPL249C | GYP5 |
| YER078C | ICP55 | YGL135W | RPL1B | YLR080W | EMP46 | YPL193W | RSA1 |
| YER008C | SEC3 | YGL203C | KEX1 | YLR083C | EMP70 | YPL178W | CBC2 |
| YDR424C | DYN2 | YGL213C | SKI8 | YLR093C | NYV1 | YPL105C | SYH1 |
| YER090W | TRP2 | YGL234W | ADE5,7 | YLR114C | AVL9 | YPL096W | PNG1 |
| YEL011W | GLC3 | YGR078C | PAC10 | YLR134W | PDC5 | YPL089C | RLM1 |
| YLR034C | SMF3 | YGR105W | VMA21 | YLR375W | STP3 | YPL057C | SUR1 |
| YMR315W |  | YGR183C | QCR9 | YMR035W | IMP2 | YPL049C | DIG1 |
| YMR316C-A |  | YHL038C | CBP2 | YMR256C | COX7 | YPR003C |  |
| YML095C | RAD10 | YHL023C | NPR3 | YNL339C | YRF1-6 | YPR004C | AIM25 |
| YML108C |  | YHR010W | RPL27A | YNL330C | RPD3 | YPR124W | CTR1 |
| YML110C | COQ5 | YHR011W | DIA4 | YNL329C | PEX6 | YPR194C | OPT2 |
| YMR099C |  | YHR016C | YSC84 | YNL233W | BN14 | YJR004C | SAG1 |
| YER063W | THO1 | YHR021C | RPS27B | YNL229C | URE2 |  |  |
| YER066W | RRT13 | YIL013C | PDR11 | YNL225C | CNM67 |  |  |
| YLR246W | ERF2 | YIL162W | SUC2 | YNL218W | MGS1 |  |  |
| YLR346C |  | YJL168C | EZL1 | YNL162W | RPL41A |  |  |
| YPL217C | BMS1 | YJL046W | AIM22 | YNL145W | MFA2 |  |  |
| YPL224C | MMT2 | YKL002W | DID4 | YNL139C | THO2 |  |  |
| YCL023C |  | YKL037W | AIM26 | YNL144C |  |  |  |
| YJR087W |  | YKL048C | ELM1 | YNL098C | RAS2 |  |  |
| YJR134C | SGM1 | YKL055C | OAR1 | YNL091W | NST1 |  |  |
| YML072C | TCB3 | YKL101W | HSL1 | YNL077W | APJ1 |  |  |
| YML085C | TUB1 | YKL134C | OCT1. | YNL054W | VAC7 |  |  |
| YJL005W | CYR1 | YKL146W | AVT3 | YNL001W | DOM34 |  |  |
| YJL028W |  | YKL148C | SDH1 | YNR001C | CIT1 |  |  |

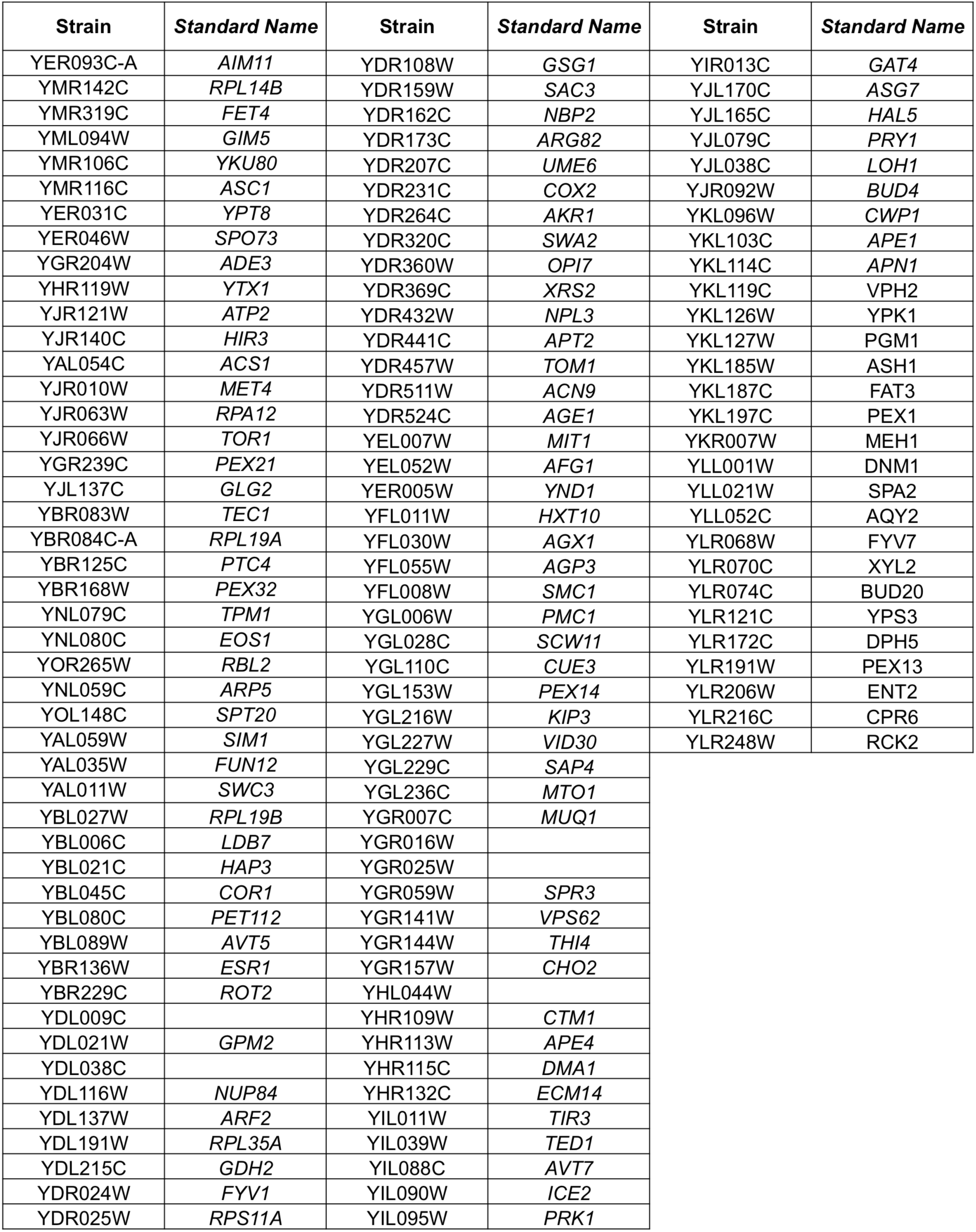
3.Desiccation Tolerant Suppressors of *tps1*Δ - 30 Day Dry - SGA

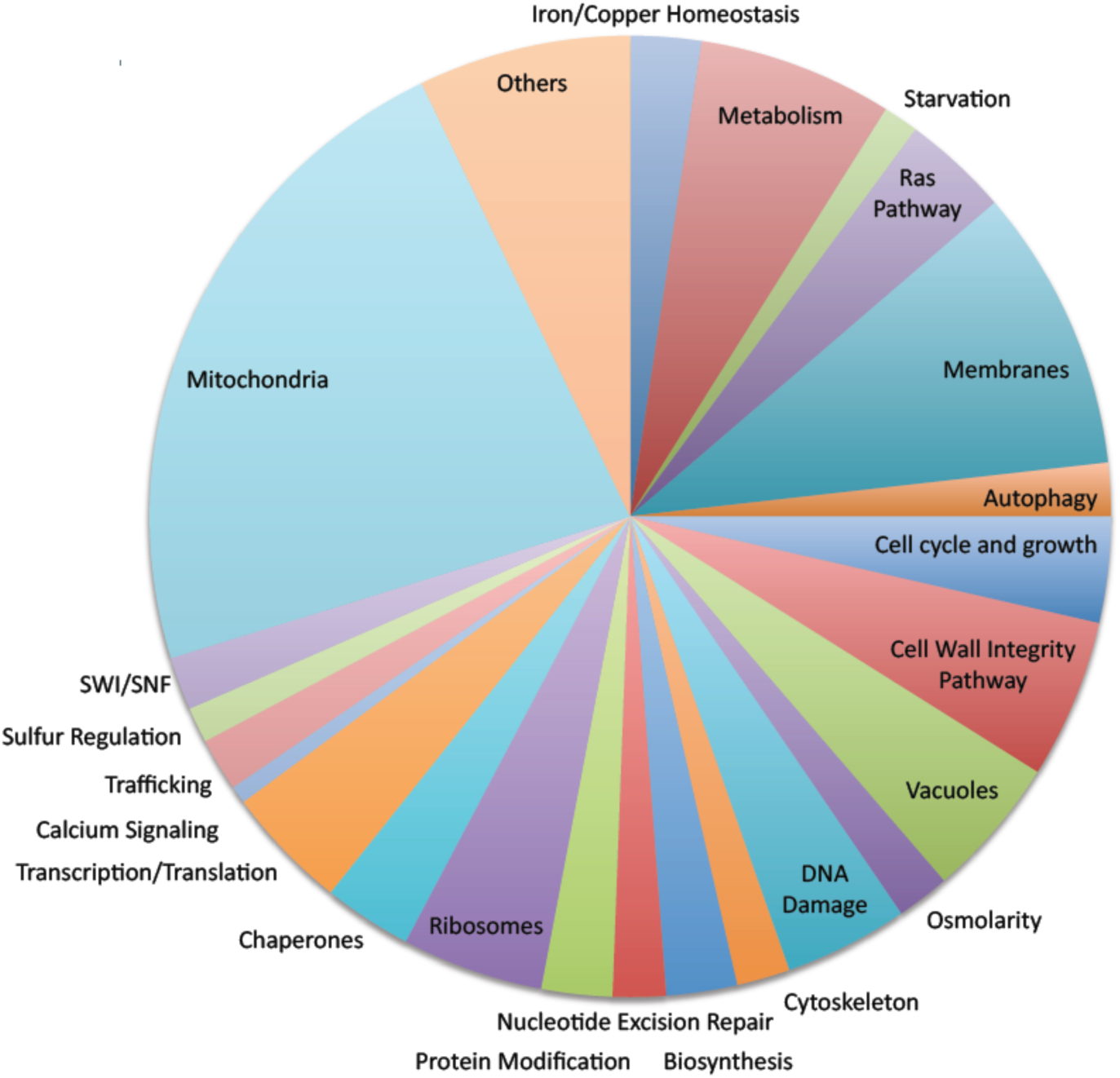
Synthetic Genetic Array Desiccation Screen – *tps1∆*. (**A**) Three different groups where identified from our SGA - *tps1∆* screen. (**1)** gene deletions that were synthetic lethal with *tps1∆*: failed to grow completely. (**2**) gene deletions that lead to desiccation sensitivity with *tps1∆*, and (**3**) gene deletions that allowed *tps1∆* to grow after 30 days of drying: suppressors. List of genes attached as 1-3. Breakdown of desiccation sensitive *tps1Δ* double mutants into categories based on cellular function. Each desiccation sensitive double mutant was placed into a category based on their cellular function (GO).

**Table S2.**
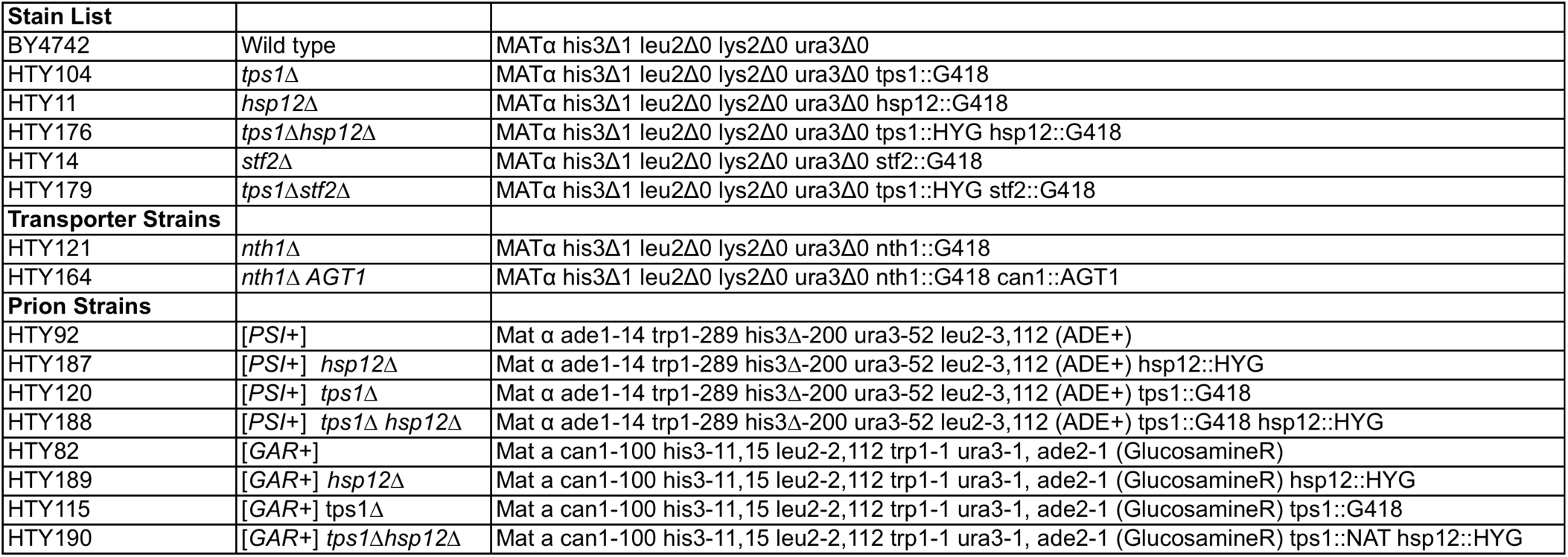
Strain List.

| Stain List |  |  |
| --- | --- | --- |
| BY4742 | Wild type | MATα his3Δ1 leu2Δ0 lys2Δ0 ura3Δ0 |
| HTY104 | *tps1*Δ | MATα his3Δ1 leu2Δ0 lys2Δ0 ura3Δ0 tps1::G418 |
| HTY11 | *hsp12*Δ | MATα his3Δ1 leu2Δ0 lys2Δ0 ura3Δ0 hsp12::G418 |
| HTY176 | *tps1*Δ*hsp12*Δ | MATα his3Δ1 leu2Δ0 lys2Δ0 ura3Δ0 tps1::HYG hsp12::G418 |
| HTY14 | *stf2*Δ | MATα his3Δ1 leu2Δ0 lys2Δ0 ura3Δ0 stf2::G418 |
| HTY179 | *tps1*Δ*stf2*Δ | MATα his3Δ1 leu2Δ0 lys2Δ0 ura3Δ0 tps1::HYG stf2::G418 |
| **Transporter Strains** |  |  |
| HTY121 | *nth1*Δ | MATα his3Δ1 leu2Δ0 lys2Δ0 ura3Δ0 nth1::G418 |
| HTY164 | *nth1*Δ *AGT1* | MATα his3Δ1 leu2Δ0 lys2Δ0 ura3Δ0 nth1::G418 can1::AGT1 |
| **Prion Strains** |  |  |
| HTY92 | [PSI<sup>+</sup>] | Mat $\alpha$ ade1-14 trp1-289 his3Δ-200 ura3-52 leu2-3,112 (ADE<sup>+</sup>) |
| HTY187 | [PSI<sup>+</sup>] *hsp12*Δ | Mat $\alpha$ ade1-14 trp1-289 his3Δ-200 ura3-52 leu2-3,112 (ADE<sup>+</sup>) hsp12::HYG |
| HTY120 | [PSI<sup>+</sup>] *tps1*Δ | Mat $\alpha$ ade1-14 trp1-289 his3Δ-200 ura3-52 leu2-3,112 (ADE<sup>+</sup>) tps1::G418 |
| HTY188 | [PSI<sup>+</sup>] *tps1*Δ *hsp12*Δ | Mat $\alpha$ ade1-14 trp1-289 his3Δ-200 ura3-52 leu2-3,112 (ADE<sup>+</sup>) tps1::G418 hsp12::HYG |
| HTY82 | [GAR<sup>+</sup>] | Mat a can1-100 his3-11,15 leu2-2,112 trp1-1 ura3-1, ade2-1 (GlucosamineR) |
| HTY189 | [GAR<sup>+</sup>] *hsp12*Δ | Mat a can1-100 his3-11,15 leu2-2,112 trp1-1 ura3-1, ade2-1 (GlucosamineR) hsp12::HYG |
| HTY115 | [GAR<sup>+</sup>] *tps1*Δ | Mat a can1-100 his3-11,15 leu2-2,112 trp1-1 ura3-1, ade2-1 (GlucosamineR) tps1::G418 |
| HTY190 | [GAR<sup>+</sup>] *tps1*Δ*hsp12*Δ | Mat a can1-100 his3-11,15 leu2-2,112 trp1-1 ura3-1, ade2-1 (GlucosamineR) tps1::NAT hsp12::HYG |

